# Machine-guided cell-fate engineering

**DOI:** 10.1101/2022.10.14.512279

**Authors:** Evan Appleton, Jenhan Tao, Greg Fonseca, Songlei Liu, Christopher Glass, George Church

## Abstract

The creation of induced pluripotent stem cells (iPSCs) has enabled scientists to explore the derivation of many types of cells. While there are diverse general approaches for cell-fate engineering, one of the fastest and most efficient approaches is transcription factor (TF) over-expression. However, finding the right combination of TFs to over-express to differentiate iPSCs directly into other cell-types is a difficult task. Here were describe a machine-learning (ML) pipeline, called *CellCartographer*, for using chromatin accessibility data to design multiplex TF pooled-screens for cell type conversions. We validate this method by differentiating iPSCs into twelve diverse cell types at low efficiency in preliminary screens and then iteratively refining our TF combinations to achieve high efficiency differentiation for six of these cell types in < 6 days. Finally, we functionally characterized engineered iPSC-derived cytotoxic T-cells (iCytoT), regulatory T-cells (iTReg), type II astrocytes (iAstII), and hepatocytes (iHep) to validate functionally accurate differentiation.

## 1 Introduction

It is not know exactly how many human cell types exist, but current estimates put the number in the hundreds [1], all originating from a single ‘totipotent’ embryonic stem cell. Since the creation of induced pluripotent stem cells (iPSCs) [2], scientists have been trying to recreate differentiation of iPSCs into all of these other types of cells and combine them into tissues or tissue-like structures (a.k.a. ‘cell-fate engineering’). This goal seems feasible given that it has been generally accepted that iPSCs are functionally identical to embryonic stem cells (ESCs) [3].

To perform cell-fate engineering, a litany of approaches have been employed that fall into three general categories: (1) application of growth factors into media in either 2D or 3D cell culture [10, 11], (2) modifications to cell matrix and plate surface conditions [12], and (3) over-expression of transcription factors (TFs) [13]. Generally speaking, the first two categories of approaches have been effective in differentiating many different cell types simultaneously — this makes sense because the general idea is to recapitulate aspects of natural development *in vitro*, where many cell types would differentiate in unison with each other. The drawbacks of these first two approaches are threefold: first, these protocols typically take a long time (often many weeks); second, the efficiency in converting to a single type of cell is often poor; and third, reproducibility across these experiments remains a large challenge. Because TF-based approaches directly manipulate the epigenetic landscape of individual cells [14, 15, 16, 8], they have proved to address these three issues to a great extent.

While TF-based approaches have been fruitful, the task of identifying the correct TFs for a fast, efficient, and robust cell conversion remains a challenging problem. There are two general ways to go about this research process: (1) an exhaustive literature search for potentially relevant transcription factors for a desired cell type and identify successful combinations via trial-and-error or (2) to use computational tools to predict TFs. While iPSCs were created through a systematic version of the former [2], this process does not scale — it is very laborious, requires deep expertise of the cell types being converted, and can only account for previously studied TFs associated with specific cell types. The latter approach has been successful in recent years [17, 18, 19, 20] and can be used as a more general approach in minimizing time required to identify effective conversion factors. While these tools have demonstrated some predictive power, they have key limitations: (1) they cannot account for experimental details such as DNA copy count, clonality (i.e. polyclonal v. monoclonal cell lines), expression method, or cell culture conditions; (2) they generally only provide a single combination of TFs for a cell-type conversion that cannot be iteratively revised; and (3) the experimental validation of a vast majority of the new outputs from these remain untested and are optimized towards very small sets of TF-combinations validated in the literature. Therefore, most tools are not geared towards finding novel TF combinations for direct trans-differentiation that may be faster and more efficient than prior reporting (**Supplementary Table 1**) [21, 17, 22, 23, 24, 9, 25, 26, 27, 28, 29, 30, 31] and (4) only one other known tool explicitly attempts to select combinations for maximal experimental differentiation efficiency [22] and no other known tool aims to maximize the speed of these differentiations. Moreover, while iterative, machine-learning (ML)-driven screening pipelines have yielded impressive results in various areas of molecular biology to date [32, 33, 34, 35], currently no tools use iterative, ML-driven screening platforms to discover novel TF combos for extremely fast cell-fate engineering.

To address these gaps, we built an epigenetics-based, ML-driven pooled screening tool for engineering cell-fate, called *CellCartographer. CellCartographer* uses next generation sequencing based readouts of chromatin accessibility (eg. DNase-seq, ATAC-seq, ChIP-seq) and transcription (RNA-seq) to predict TFs to be correlated with cell-type identity. Using the predictions made by *CellCartographer*, we can define multiplex pooled-screens of TFs for over-expression, which allows us to explore many experimental variables such as variable stable expression quantities, genomic integration copy count and location, and culture conditions with the option to add more nuance depending on experimental conditions. Furthermore, we can implicitly select TF over-expression combinations based on speed and efficiency depending on the differentiation and screening conditions. *CellCartographer* gives outputs agnostic to starting cell type because it has been demonstrated that the same (or similar) TF set can be used to differentiate cells from a variety of originating cell types [22] and because the iterative engineering process from this starting *in silico* screen should be able to accommodate for these differences. We demonstrate how the *CellCartographer* predictions are sufficient for differentiating small sub-populations of cell-surface marker-positive cells for twelve target cell-type samples from all three germ layers (resulting in the exploration of up to 10 million TF combinations for these twelve types). We then show how we can use bulk-RNA sequencing to refine the original TF predictions and zoom in on minimal TF combo sets to differentiate stem cells for six cell types from all three germ layers. Once a sufficiently-high percentage of polyclonal cell line differentiation was created, we showed that isolating clones from these populations results in the creation of high-performance clonal lines with extremely high differentiation speed and efficiency. Finally, we functionally characterized robust clonal lines of differentiation-inducible iPSC lines for each of the three germ layers: regulatory T-cells (iTReg) and cytotoxic T-cells (iCytoTs) - mesoderm, hepatocytes (iHep) - endoderm, and type-II astrocytes (iAstII) - ectoderm) to validate that the cells are functional *in vitro* and molecularly accurate. We were able to differentiate four cell types using novel combinations of TFs in as little as 6 days. Importantly, our unprecentdented derivation of iTRegs and iCytoTs directly from iPSCs in simple media conditions in <6 days may considerably accelerate the investigation of T-cell biology.

## 2 Results

### 2.1 Machine learning for determining TF sub-libraries

As many TFs are controlled for activity by nuclear localization [36], RNA expression alone is not a sufficient indicator of TF activity and importance for cell identity. A stronger indicator of TF activity is occupancy of TFs at active DNA regulatory elements, which are marked by methylation and acetylation marks [37, 38]. While chromatin immunoprecipitation sequencing (ChIP-seq) [39] can be used to determine TF binding and DNA histone methylation/acetylation, performing assays for each possible pioneer factor for a cell would be infeasible. Chromatin accessibility assays such as DNAse-seq and ATAC-seq captures the super set of all transcription factor binding sites and allows for the indirect observation of TF binding through their DNA binding motifs. With the aim of minimizing resource requirements for studying a novel cell type, the *CellCartographer* model leverages chromatin accessibility data to make initial predictions of TFs for differentiating towards that cell type. After initial TF predictions are made, TF transcript levels are used to exclude TFs that are not expressed. The *CellCartographer* pipeline can leverage a variety of assays for chromatin accessibility and transcriptomics to predict a set of TFs for a target cell type, which can then be tested in a pooled screen (**Figure 1a**). To broaden the functionality of *CellCartographer*, input data can be either manually uploaded or automatically queried and downloaded from the ENCODE database [40] or GEO [41]. While we use chromatin accessibility only during our studies, additional assays that can be used to exclude inactive regulatory elements such as DNA histone acetylation/methylation and nascent RNA expression would likely improve the quality of predictions.

**Figure 1:**
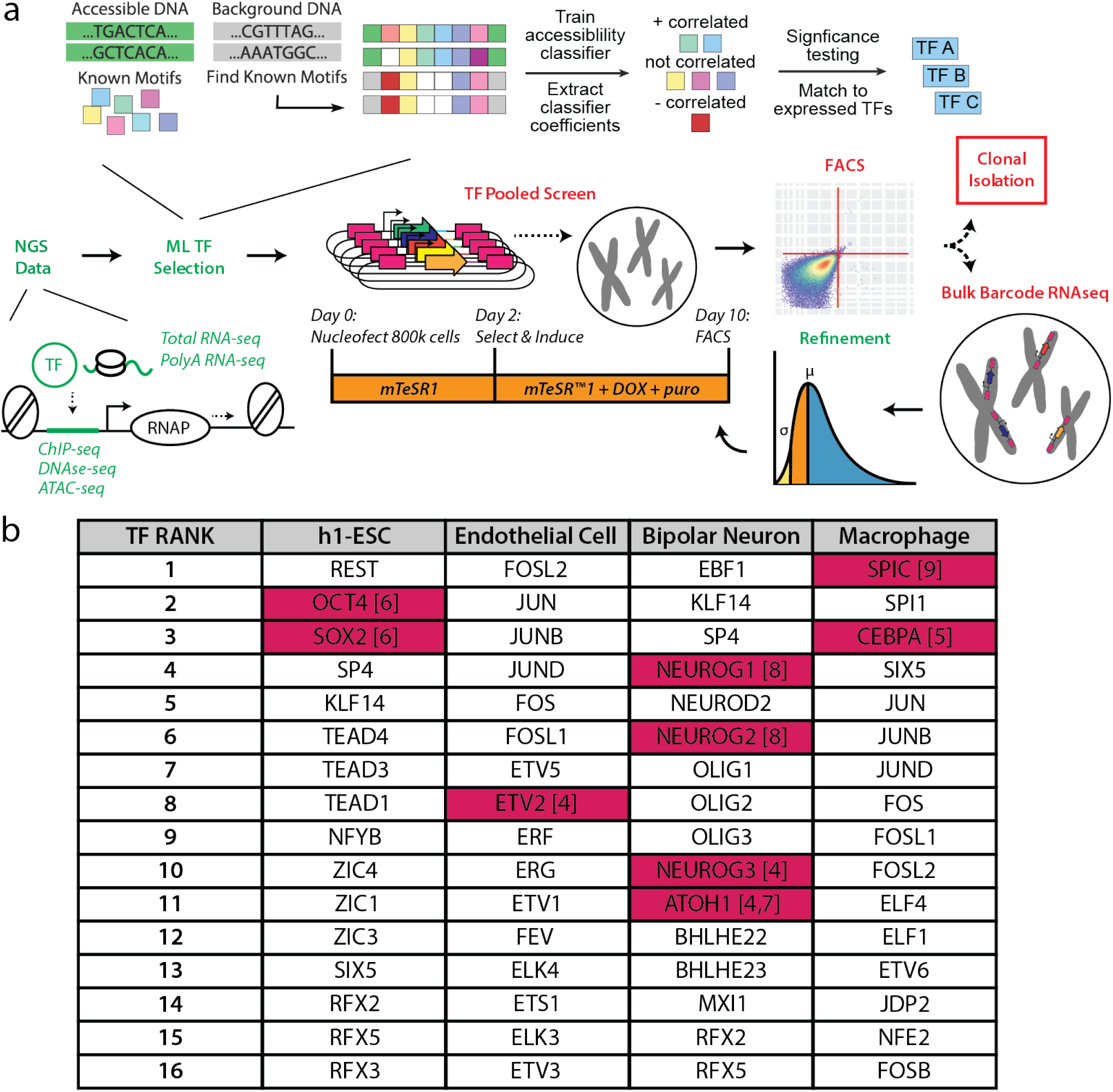
*CellCartographer* workflow. **a.** The *CellCartographer* workflow uses epigenetics, then transcriptomics NGS data (computational steps in green) to determine TF pools for iterative screening with the TFome (experimental steps in red). Iterative rounds of screening are refined with barcode sequencing and statistical analysis. Polyclonal cell lines with sufficient differentiation (>10%) undergo clonal isolation to isolate high-efficiency clones. Cells are nucleofected with barcoded TF-cassette pools that are integrated randomly into the genome where any one cell may receive some combination of these factors in either multiple copies (blue) or not at all (green/yellow), resulting in 800,000 TF combo experiments per nucleofection. The distribution of TFs that are delivered to cells’ nuclei is approximated by a Poison distribution that can be statistically evaluated to refine screens to TFs **b.** *In silico* validation of screening lists — for four cell types with previously validated TF-overexpression differentiation factors [4, 5, 6, 7, 8, 9], our model accurately re-identifies these factors (magenta) in the top TFs that would be put into a screen.

Since the number of TFs in the TFome (1732) with characterized binding sites [42] (891), yields 2^891^ possible outcomes (**Figure 1a**), a full library screen is intractable. In a full library screen, the chance of observing a correct combination of TFs that differentiate a target cell type with 10^6^ starting cells would be unlikely (on the order of 1 in 10^167^). And so, we reasoned that the number of starting cells and the number of possible combinations formed from the set of transfected TFs should be similar (i.e. 2^*n*^*T F s* m_*cells*_). In our case, we nucleofected 10^6^ cells per experiment and the screening pools contained approximately 16 plasmids containing integratable TF-over expression cassettes driven by a doxycycline-inducible promoter (**Figure 1a**). Each TF cassette is integrated randomly within each cell from zero to *n* times, allowing us to explore a large parameter space of DNA integration location and resulting expression amounts of each TF in combination.

In order to identify TFs of interest for each cell type, we begin by learning a relationship between TF motifs and chromatin accessibility. Specifically, we train a logistic regression classifier model to distinguish between open chromatin regions and a set of background genomic loci, matched for GC content, using known DNA TF binding motifs drawn from the JASPAR database (**Figure 1a**) [43]. By training a model using all motifs, we can model the cooperative binding of lineage determining TFs [9, 44]. Given that we want to select a small number of TFs, we use LASSO regularization when training the model. One dvantage of linear models in comparison to more complex models such as deep neural networks, is greater interpretability. By examining the sign of the model coefficients, we can determine whether the presence of a motif is negatively or positively correlated with open chromatin. We exclude all TF motifs that have a negative coefficient and are negatively correlated to open chromatin. Correlated features, in our case similar TF motifs, can result in multiple-collinearity and unstable model coefficient values. We mitigate multiple-collienarity by first using a non-redundant set of motifs [45]; additionally, we train an ensemble of models across five cross validation splits and use the mean result across the ensemble. To further increase the stringency of our results, we determine the significance of each TF motif using the likelihood ratio test, which is an *in silico* analog of a mutagenesis experiment. In the likelihood ratio test, the performance of a model trained on all motifs is compared to a model trained on all but one motif. We can identify and exclude constitutively active TFs pooling results from across several cell types; we rescale (z-score normalization) the coefficients of models for all cell types (**Supplementary Figure 1**) and remove all motifs that has a mean absolute z-score greater than 2.5 (**Supplementary Figure 2**). As many TFs share DNA-binding motifs [46], we then use transcriptomics data to identify which TFs are expressed in a given cell type; we select the most significant motifs that are positively correlated with binding and the top 16 corresponding genes that have RNA expression. Our procedure for selecting TFs for testing is outlined in (**Figure 1b**) and (**Supplementart Figure 3**). Using publicly available DNase-seq data from ENCODE, we applied our approach on several cell types with simple known combinations of one to two lineage determining TFs and confirmed that these TFs appear in the top TFs predicted by *CellCartographer* (**Figure 1b**).

To computationally validate our model on a larger scale, we applied *CellCartographer* to 34 primary cells types and 29 tissue types. We found that each TF DNA binding motifs strongly correlated with chromatin accessibility had different behaviors in each cell type and tissue type (**Figure 2c, Supplementary Figure 5c**). Given that related cell types have similar transcriptional profiles (**Figure 2a, Supplementary Figure 5a**), we reasoned that they may also have similar TFs correlated with open chromatin that drive transcriptional profiles. To visualize the similarity between transcriptional profiles, we calculated the pairwise Pearson correlation between the gene expression values of each cell type (log RPKM values) and used multidimensional scaling to embed each cell type in way that respects the pairwise similarity between cell types; using the Spearman correlation model coefficient for each TF, we can also visualize the similarity of TF motifs correlated with open chromatin. We observe that cell types that group together when considering similarities in transcriptional profiles such as adaptive immune cells (eg. B-cells and T-cells) and progenitor cell types (H1-hESC, GM23338, and neural stem progenitor) tend to look similar from the perspective of TF motifs correlated with open chromatin (**Figure 2b, Supplementary Figure 5b)**.

**Figure 2:**
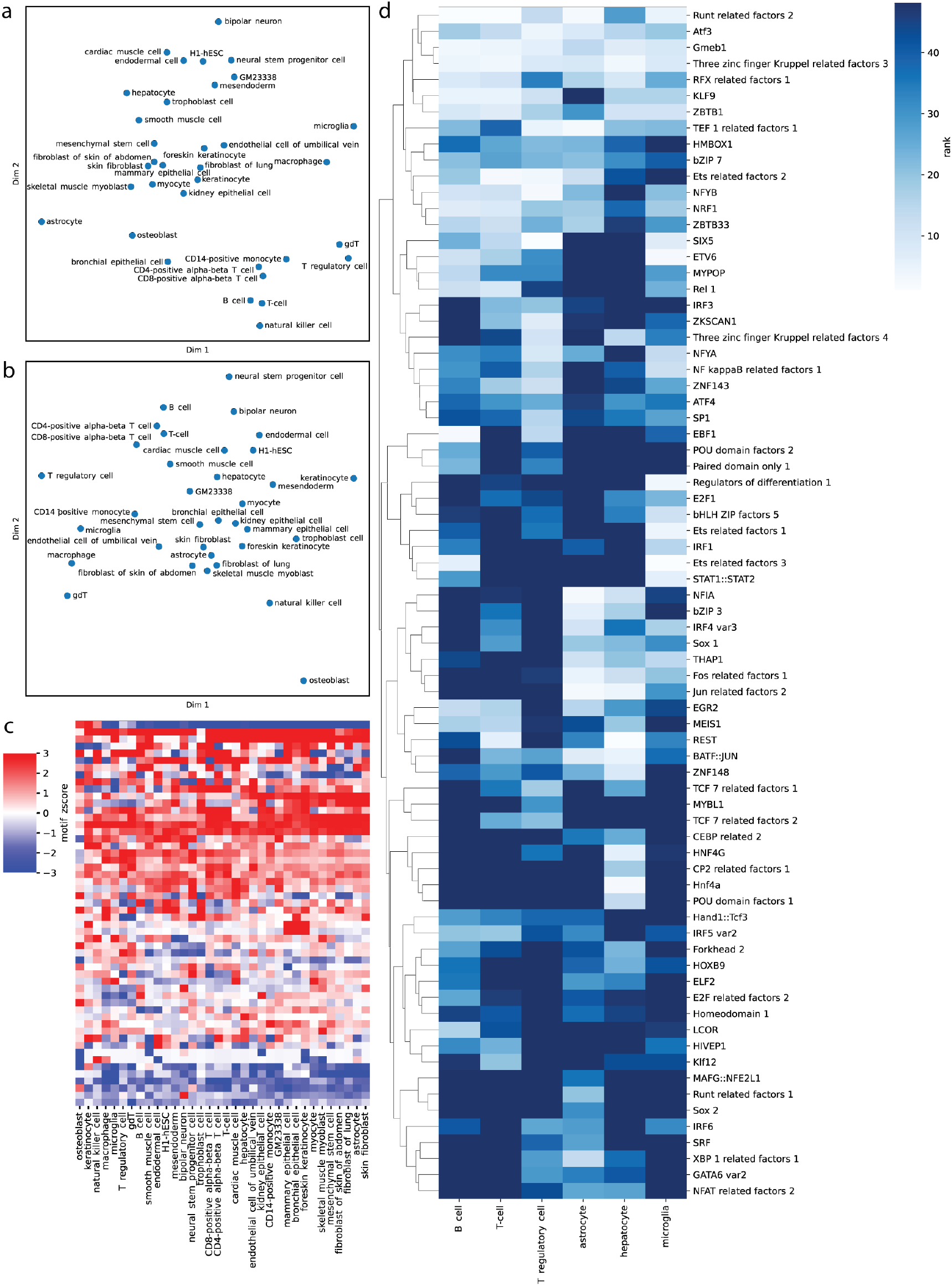
Computational analysis of 34 cell types with *CellCartographer*. **a.** Multidimensional scaling of the similarity in gene expression between different cell types. **b.** Multidimensional scaling of the similarity in TFs correlated with open chromatin. **c.** Motifs correlated (red) and anti-correlated (blue) with open chromatin vary across 34 cell types analyzed. **d.** Highly ranked motifs correlated with open chromatin for cell types derived from yolk sac (microglia), endoderm (hepatocyte), mesoderm (B-cell, T-cell, regulatory T-cell), and ectoderm (astrocyte)

### 2.2 Primary pooled TF screens for differentiation

To demonstrate that our pooled screening method could be generally applied to any cell type of interest, we identified cell types from each human germ layer and screened TFs combinations to identify populations of cells that came up positive for canonical markers. Specifically, we generated TF pools for: Mesoderm — T-cells (subtypes cytotoxic, delta-gamma, and regulatory), B-cells, macrophages, epithelial cells (subtypes kidney, bronchial, and mammary), and osteoblasts; Endoderm — hepatocytes; Ectoderm — type II astrocytes; and Yolk Sac — microglia. For each cell type, we designed two TF pools for each cell type using *CellCartographer* — one pool containing TFs with expression level *≥* 1 RPKM and another containing TFs with expression level *≥* 4 RPKM (**Supplementary Tables 1-12**). We then prepared mixed DNA pools of equal concentration of each TF and nucleofected and screened iPSCs for differentiation (**Figure 1d**). We found that the percentage of cells appearing positive in most cases was very small, but ranged from 0.05% (Regulatory T-cells) to 17.64% (B-cells), although in almost all cases, the positive population was <1% (**Figure 3, Supplementary Figure 7**). Thus it appeared that all samples yielded at least a small population of differentiated cells that can be sequenced to determine which TFs from the TFome were present.

From this set of diverse screened cell types, we decided to iteratively refine a set of six that had high clinical relevance — cytotoxic T-cells, Regulatory T-cells, B-cells, hepatocytes, type II astrocytes, and microglia. A comparison of the top motifs positively correlated with open chromatin for these six celltypes is shown in **Figure 2d**; the screening pools for each of these six celltypes are shown in **Supplementary Table 1-6**. It should be noted that at this step, the selection of specific surface markers biases the downstream analysis and refinement. For example, although TF pools for astrocytes were determined from data based on generic astrocytes (type I or type II), our selection of A2B5 as a surface marker in combination with CD44 selected specifically for type II astrocytes. In the case of the epithelial sub-types, there was some uncertainty of the ideal cell surface markers to use since CD24 was unexpectedly present in the stem cells and stem cells are partially epithelial in quality and express EpCAM [47] to a slightly lesser degree than differentiated epithelial types. Nonetheless, from the pooled screens we were able to sort at least 1000 double-positive cells from each large population for bulk RNA-sequencing. We lysed the sorted cells, prepared sequencing libraries, and amplified the barcoded regions of the TFome cassettes to tell us the relative abundance of TFome cassettes in the double-positive cells (**Figure 3**). We found that the distributions for each cell type had some variability, but that in general, each cell type had TFs that were represented in the positive population more than others. In fact, only one of the six cell types (cytotoxic T-cells) had all TFs show up in sequencing at least once.

**Figure 3:**
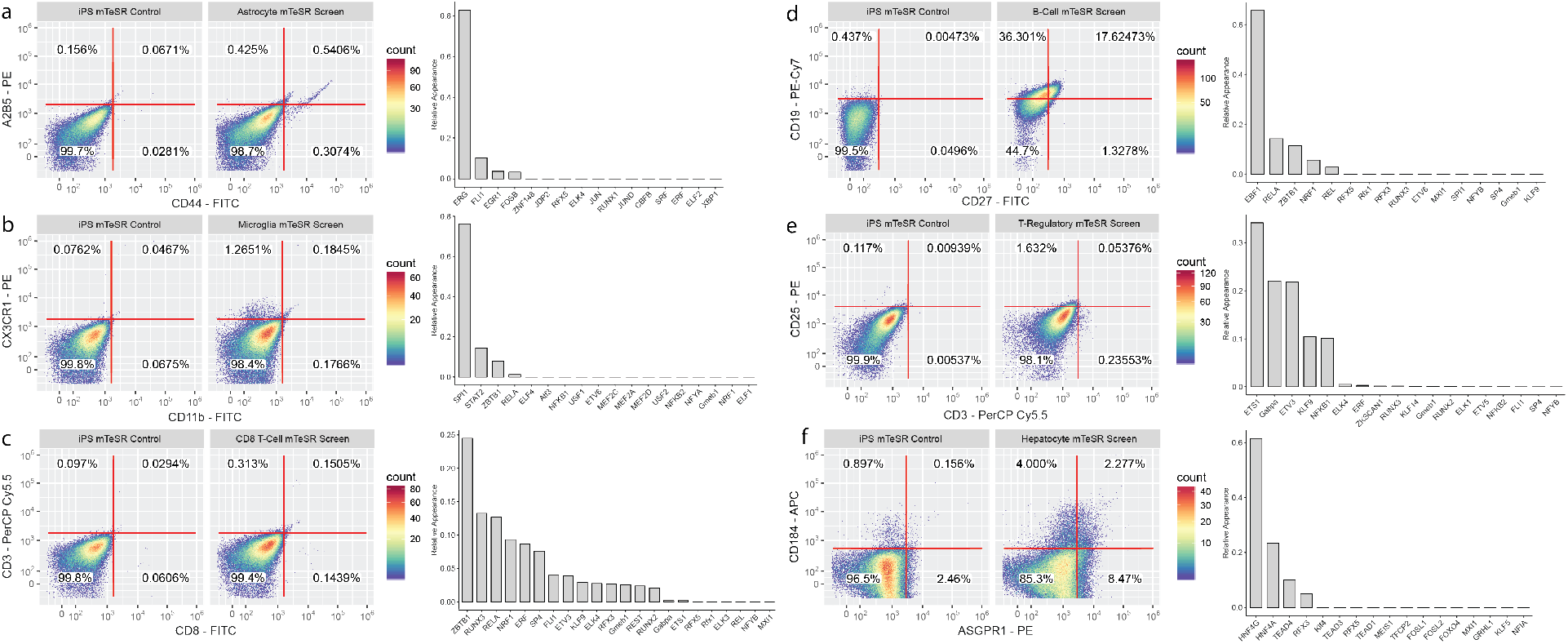
Primary pooled screens for cell types originating from each germ layer. For each cell type, a negative antibody stain for iPSCs without TFs (LEFT), the cell population with induced TFs (MIDDLE), and the barcoded TF appearance frequency in the transcriptome of double-positive cell populations (RIGHT) is shown. **a.** Type II Astrocytes (ectoderm) **b.** Microglia (yolk sac) **c.** CD8-positive T-cells (mesoderm) **d.** B-cells (mesoderm). **e.** Regulatory T-cells (mesoderm) **f.** Hepatocytes (endoderm).

### Iterative pooled TF screening and clonal isolation

Using the barcode frequencies, we calculated 3 refined TF pools for each cell type: All TFs that appear in sequencing, TFs that appear greater than average, and TFs that appear one standard deviation or more than average (**Figure 4a-d**. Using the refined TF pools, we performed a second round of differentiation. Given that this round of screening generally limited TF pools to <5 TFs per pool, we built stable cell lines for additional testing and refinement. iPSCs were nucleofected as before, but we selected and stabilized the cell lines before screening differentiation in different settings. Specifically, given the stability of the constructed cell lines (i.e. less cell death), we opted to test them for only six days, and also decided to test their performance in target-cell-type growth medium in addition to stem cell medium (**Supplementary Figure 8**).

**Figure 4:**
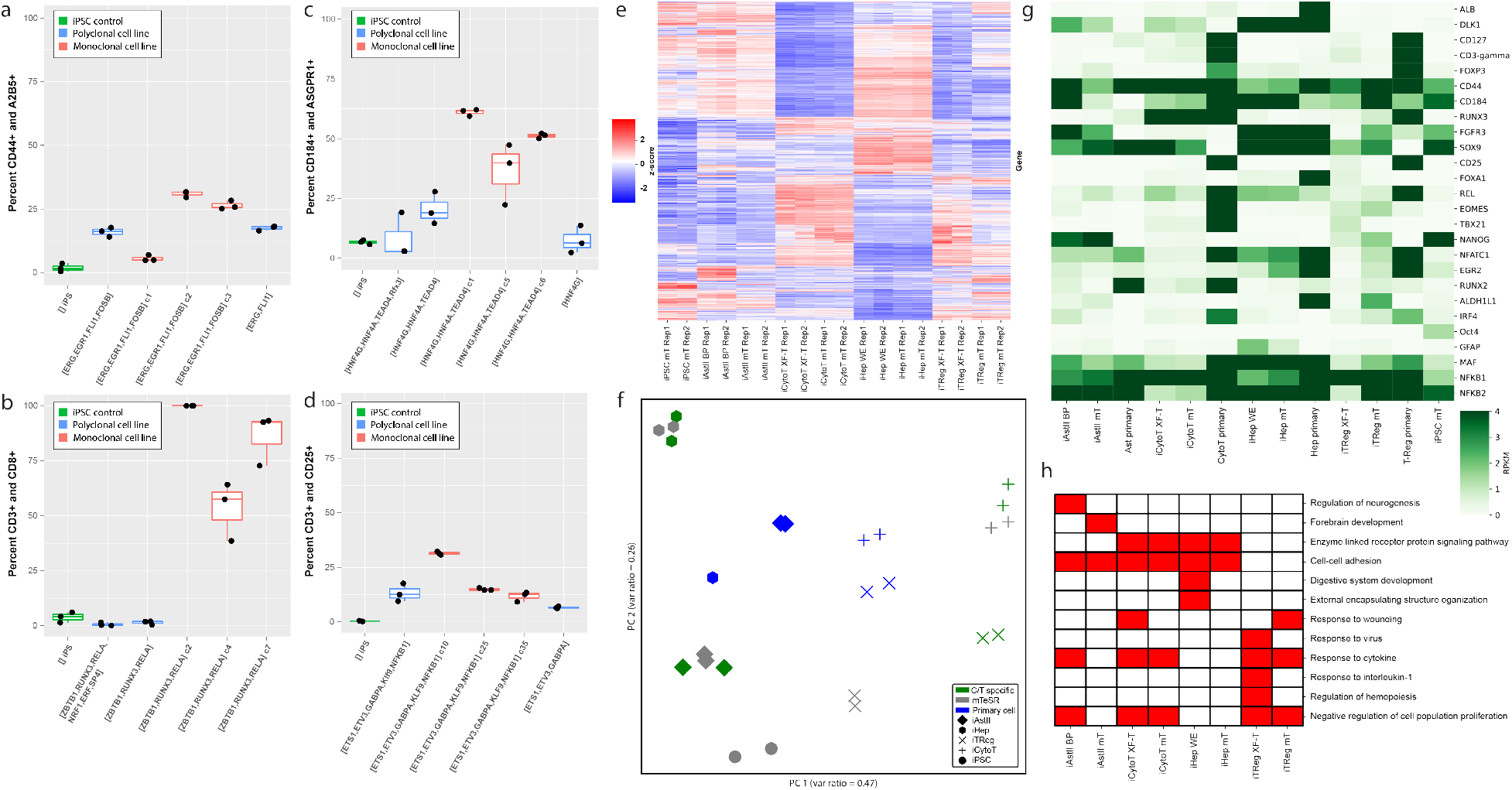
Iteratively engineered poly-clonal and mono-clonal cell lines. (**a-d**) For each cell type, we show percent double-positive for FACS analysis of canonical markers for non-clonal (turquoise) and mono-clonal (red) cell lines mono-clonal (red) cell lines, and an iPSC + media control (green) differentiated for six days in cell-type-specific media + DOX for **a.** Type II Astrocytes (iAstIIs) **b.** Cytotoxic T-cells (iCytoTs) **c.** Hepatocytes (iHeps) and **d.** Regulatory T-cells (iTRegs). **e.** Differential gene expression (quantified by Z-score) for all genes for two replicates of each differentiated cell type in both media conditions. **f.** Principal component analysis of all genes for each cell type in each media condition and a primary cell control. **g.** Differential gene expression (quantified by RPKM) for key marker genes across target cell types and iPSCs. **h.** Metascape [48] analysis of gene enrichment of high-efficiency clones for genes that were upregulated in these lines and differentiation conditions compared to iPSCs. Analysis of select highly-significant GO Terms from TOP 50 for each differentiated cell type and condition is shown (-log10(P) *≥* 3).

In this round, we found broad improvement in differentiation percentage across all six cell types (**Figure 4a-d, Supplementary Figure 11**). While B-cells already had a considerably high differentiation percentage in the primary screening round (17.6%), it improved to an average of greater than 50%. For all other five cell types, the refined lines appeared to improve in differentiation percentage dramatically compared to the populations seen in the primary screen. However, since these populations have mixed identity, it is likely that many of these cells were still only partially differentiated. When we examined the number of cells that were positive for just one (or both) markers, all cell types improved differentiation rates compared to the primary screens (**Supplementary Figure 10**). When we examined differentiation percentage (both partial and total) in target-cell-type growth media, we saw even more near-complete differentiation of these cell lines (**Supplementary Figure 8,12**). While it was clear that the growth medium is a contributor to differentiation efficiency, the TFs were the major driver of differentiation for all cell types.

Given that our cell lines were clearly making progress towards robust differentiation, but in a limited capacity, we reasoned that perhaps many micro-scale experimental details could be to blame — for example, perhaps cell-cell communication from non-differentiating cells in the population was the issue, or perhaps the details of how many TF cassettes were integrated and in what location was very important. Since we use PiggyBac integrase that integrates variable copies of TF over-expression cassettes in random genomic locations, we hypothesized that perhaps some cells in the cell line population are holding back the rest of the population, and that isolating monoclonal cell lines could improve our differentiation efficiency. Ergo, we sorted random single cells in the population to form monoclonal lines and characterized them. To our satisfaction, for CD8 T-cells, microglia, astrocytes, and hepatocytes, this solved the problem — several clones of each were able to dramatically outcompete the mixed population in differentiation efficiency in all of the aforementioned differentiation conditions (**Figure 4a-d, Supplementary Figure 11**).

After differentiation of high-performance clones, we performed RNA-sequencing to validate that our clones were generally reflective of target cell types at a molecular level in addition to surface markers. We found that across all genes, our differentiated cells clustered well by cell type in both media conditions (**Figure 4e**). Specifically, it was important to see that the molecular characteristics of the T-cell subtypes were in general agreement and were significantly different from all other types. As expected, since these cell types were all from different germ layers (except the T-cell subtypes), the expression profiles were dramatically different across differentiated cell types. This was further reflected in principal component analysis (**Figure 4f**) - we observed that our differentiated cell types generally clustered very tightly across both media conditions and that they clustered somewhat well with primary cell types. The clustering of cell types across variable media reinforces that TF over-expression is a more dominant factor than the different media conditions. Next, when we zoom in on key canonical markers for our differentiated cells, they once again cluster as expected and generally show upregulation of expected markers (**Figure 4g**). In the case of iAstIIs and iTRegs, there were some interesting marked difference of key factors across media conditions, suggesting that media formulation may play a key role in the final condition and function of these cells. Finally, when we analyze the complete sets of significantly up-regulated genes (P < 0.1) for our high-efficiency clonal lines compared to iPSCs with Metascape [48], we see enrichment of GO terms that is supportive of cell-type specific features (**Figure 4h**).

### Functional characterization of differentiated cells

Finally, after refinement of our differentiating cell lines and molecular validation of their identities, we wanted to validate that the cells also functionally perform their intended function for down-stream clinical applications. To this end, we opted to focus on at least one cell type from each germ layer - regulatory T-cells (iTRegs), cytotoxic T-cells (iCD8s), type II astrocytes (iAstIIs) and hepatocytes (iHeps). To functionally characterize these cell types, we performed *in vitro* assays based on biological function (**Figure 5**).

**Figure 5:**
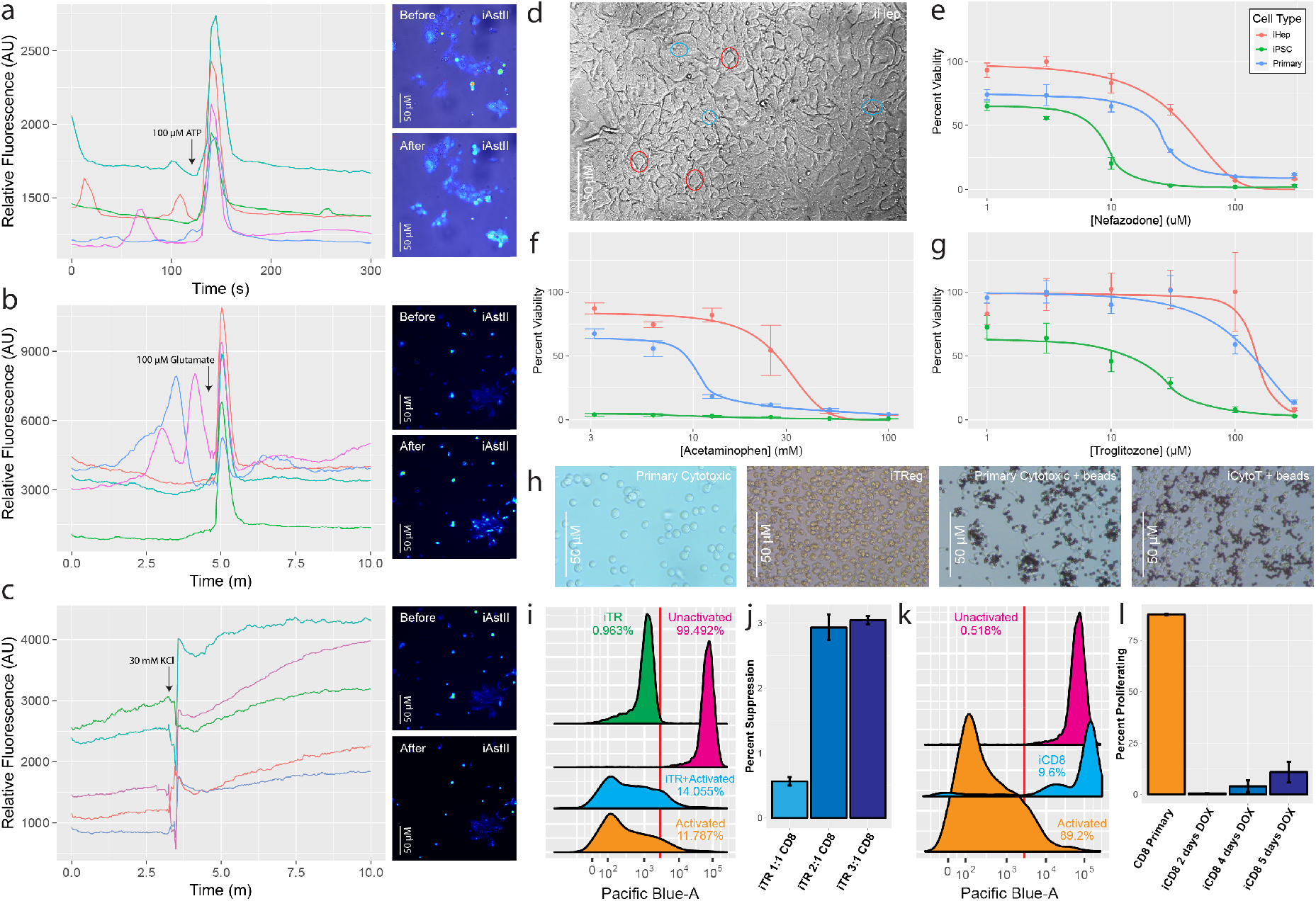
Functional validation of iAstIIs, iHeps, iCytoTs, and iTRegs. **a-c.** Stimulation of Type II astrocytes over 10 min with small molecules with **a.** 100*µ*M ATP **b.** 100*µ*M glutamate, and **c.** 30mM KCl. (LEFT) Relative fluorescence of six individual astrocytes. Astrocyte cell population shown before (TOP) and after (BOTTOM) addition of small molecule. **d.** Phase-contrasted BF image of induced iHeps prior to hepatotoxicity testing. Key features in select cells such as multiple nuclei (blue circles) and approximately cuboidal shape (red circles) are called out. iHeps, primary hepatocytes and iPCs titrated with **e.** Nefazodone, **f.** Acetaminophen, and **g.** Troglitazone for 24h and assayed for percent viability (survival rate normalized to each cell type without toxins applied). **h.** Brightfield imaging of T-cell populations (LEFT to RIGHT): Primary CD8 T-cells, iTRegs, Primary CD8 T-cells + activation beads, iCytoTs + activation beads. **i.** Suppression assay for iTRegs co-cultured with activated primary CD8 T-cells **j.** Calculated percent suppression with titrated dosing of iTRegs in suppression assay. Primary T-cells have been shown to suppress in the range of 20/30/40% respectively [49]. **k.** Activation assay for iCytoTs **l.** Percent of proliferating primary CD8 T-cells and iCD8 cells post-activation.

For the iAstIIs, we validated that the morphology was correct and that they were stimulated as expected by certain standard small molecules [50] (**Figure 5a-c**). We observed that at standard concentrations of small-molecules of three classes (glutamate - neurotrasmitter, ATP - nucleotide, and KCl - ionic), many plated astrocytes were stimulated. We observed strong increases of relative Fluo-4 fluoresence immediately after induction for individual astrocytes that were both inactive before stimulation and active at times before stimulation. Furthermore, while glutamate and KCl should stimulate both astrocytes and other neuronal types, only astrocytes are stimulated by ATP, confirming that the cells we assayed both had correct astrocyte morphology and exclusive functionality.

For the iHeps, we validated the morphology (**Figure 5d**) and compared their viability compared to primary hepatocytes and undifferentiated cells when exposed to hepatotoxins for 24hrs [51, 52]. We observed that our iHeps had highly similar viability to primary hepatocytes after being exposed to Nefazodone (**Figure 5e**), Acetaminophen (**Figure 5f**), and Troglitozone (**Figure 5g**), and demonstrated significantly higher viability compared to undifferentiated iPSCs.

iTregs were validated by demonstrating that the cells inhibited the expansion of responder T-cells [53]. Before this step, we confirmed that our iTRegs had size and morphology approximately the same as primary cytotoxic responder cells (**Figure 5h**). While the size and shape were generally consistent, with both iTRegs and iCytoTs, the primary responder T-cells took on an elongated shape when stimulated, while our iCytoTs did not clearly show this morphological change to stimulus. Responder T-cells were stimulated to activate with IL-2 and CD3+CD28+ beads for three days. After this activation step, responder T-cells were labeled with a fluorescent dye and co-cultured with iTregs in variable quantities. After 11 days, fluorescence was recorded to validate that the addition of more iTregs resulted in reduced responder T-cell proliferation (**Figure 5i,j**). We observed some reduction in responder T-cell proliferation as we increased the number of iTRegs, albeit modestly compared to prior results with primary regulatory T-cells [49]. Finally, to validate the iCytoTs, we activated them with the same bead-based method used in the the regulatory T-cell assay and examined their morphology and interaction with the activator beads (**Figure 5h**) and then recorded proliferation. We found that as with the iTRegs, the proliferation was modest, but increased by the number of days the iCytoTs were induced from stem cells prior to the initiation of the assay (**Figure 5k,l**).

## 3 Discussion

In summary, we have described how the *CellCartographer* tool and pipeline can guide and refine cell-fate engineering with machine learning and synthetic TF-cassettes from the human TFome. We demonstrated that the primary TF pools for differentiating iPSCs into a diverse set of cell types yields a small population of positive cells for each of the tested types. We then went on to focus on six cell types from each germ layer to show how we can use NGS data from partially-engineered cell lines with *CellCartographer* to engineer high-efficiency differentiation-inducible cell lines. Finally, we isolated high-performance clones for four cell types and functionally characterized at least one cell type from each germ layer to validate that our engineered cell lines were functionally accurate *in vitro*.

While *CellCartographer* is not the first software to identify TFs for cell-fate engineering, it presents an advance in three main areas from a software perspective. First, it leverages a machine-learning driven, iterative screening pipeline by making TF predictions using epigenetics data and enables an iterative pipeline for refining engineered cell lines. We hypothesize that as sequencing technologies continue to improve and more data is generated, *CellCartographer*’s predictions should only improve. Second, *CellCartographer* has a very minimal requirement for producing useful TF pools — it does not require re-training large models for additional cell types, which can prove useful for engineering cell lines for differentiation into exotic cell types with little data available. Furthermore, we were able to successfully engineer iTRegs using TFs determined from *Mus Musculus* data since that was the only epigenetic NGS data available for this cell type, meaning calculations of factors can work cross-species. Finally, the pooled screening philosophy of *CellCartographer*, allows biologists to explore and debug many experimental variables that are generally invisible to software tools — namely synthetic DNA genomic integration location, copy count, and cell culture conditions and their resulting differentiation speed and efficiency. Pooled screening and paired ML analysis allows us to screen out these issues. Furthermore, while we use the starting predictions from *CellCartographer* to iteratively refine our cell lines in this study, all of the down-stream tools are compatible with starting predictions from other tools (i.e. another tool could provide the starting prediction and *CellCartographer* and the TFome can still be used downstream), meaning *CellCartographer* can be used to compliment other existing tools that generate complete lists of TFs predicted to be associated with cell type [28, 24, 29] (**Supplementary Table 1.**).

This work also represents a major advance in terms of identifying four robust TF combinations for differentiation into high-value cell types relevant to therapeutics. At this time, aside from hepatocytes, there are no experimentally established TF combinations for directly differentiating stem cells into type II astrocytes, regulatory T-cells, or cytotoxic T-cells. Furthermore, we demonstrate that this differentiation can be driven in stem cell media in six days or less, meaning that the TF combinations are fast, robust, and solely to credit for the differentiation in these examples. Finally by performing additional optimizations with specialized media conditions and performing functional assays on iAstIIs, iHeps, iTregs, and iCytoTs, we show that this strategy should be robust in ultimately obtaining functional clonal cell lines of theoretically any type that can differentiate rapidly, efficiently, and robustly from iPSCs. While the functional qualities of the iAstIIs, and iHeps were more dramatic and complete, the function and viability of the induced T-cells is likely very sensitive to media conditions and could be further improved with additional optimization of growth conditions starting from the stem cell state. A clear next step from this work would be to further optimize culture conditions for these cell types to improve functionality and even perhaps to re-perform screens in these optimized medias.

In conclusion, we believe that *CellCartographer* provides a clear benefit to the field of stem cell biology and cell-line engineering. While we have already generated interesting inducibly-differentiating iPSC lines, we strongly believe that this tool can be applied immediately to aid the engineering of other stem cell lines for any number of therapeutic, diagnostic, or other commercial applications.

## 4 Methods

### DNAse-seq and ATAC-seq analysis

Adapters from sequencing reads were trimmed with Homer [9], using the command: homerTools trim -len 40. Following adapter trimming, reads were aligned using Bowtie2 [54] (with default parameters) and then converted into a Homer tag directory. We called open chromatin regions or peaks with Homer using the following findPeaks command with the following parameters -C 0 -L 0 -fdr 0.9. We then use IDR [55] to identify high confidence open chromatin regions.

### Prediction of transcription factors for cell fate engineering

For the set of open chromatin regions for each cell type, we sample from the genome an equivalent number of background peaks that has matching GC content and size. Using a set of non-redundant DNA motifs [45], which specify the frequency of each nucleotide at each position in the motif, and a background frequency (0.25 at each position), we can calculate a log odds score that indicates how well a sequence matches a motif. For each open chromatin region and background loci, we calculate the highest log odds score for each motif. We standardize the motif scores such that the mean score value is 0 and the variance is 1. Then we train a LASSO-regularized logistic regression model [56] to discriminate between open chromatin regions and background sites. We assess the importance of a motif using a log-likelihood ratio test where we compare the performance of a perturbed model where a single motif is not used as a feature during model training and the performance of the full model that is trained using all motifs. We convert the difference in likelihoods given by the two models to p-values using the chi-squared test. Model coefficients and p-values reported are the average across five randomly assigned cross-validation splits (80% training, 20% testing). As a sanity check, for each model, we measure the area under the receiver operating characteristic (ROC) curve, and ensure that the model is making non-random predictions. The model training procedure and TF selection procedure is summarized in (**Figure 1b**) and (**Supplementary Figure 3**). Data processing, model training, and statistical analysis was performed using python and the following packages: pandas [57], numpy [58], scipy [59], sklearn [60], biopython[61]. Data plotting performed using R bioconductor packages [62].

### Transcriptomics analysis

Adapters from sequencing reads were trimmed with Homer, using the command: homer-Tools trim -len 40. Following adapter trimming, reads were aligned using Bowtie2 (with default parameters) and then converted into a Homer tag directory. We used the Homer analyzeRepeats command to quantify gene expression as RPKM values. Raw read counts at each gene were used as input to DeSeq2 [63] for identifying differentially expressed genes.

### Cloning of transcription factors

Transcription factors were cloned into puromycin-resistant cassettes with flanking piggyBac transposon [SystemsBio] genomic integration regions under the control of the mammalian DOX-unducible promoter pTRET. Plasmids for each transcription factor are members of the ‘Human TFome’ library deposited on Addgene.

### Creation of cell lines and cell culture

All differentiating cell lines and differentiation screens were performed on reprogrammed PGP1 fibroblasts using the Sendai-reprogramming-factor virus. PGP1 iPS cells were expanded and nucleofected with P3 Primary cell 4D Nuceleofection kits with pulse code CB150 using 2*µ*g of total DNA for 800,000 cells (1.6 *µ*g TF pool/0.4 *µ*g SPB) [Lonza]. Cells were plated onto Matrigel-cotated plates [Corning] with ROCK-inhibitor [Millipore] and selected with puromycin [Sigma]. Stable cell lines were expanded over several passages using TrypLE [Gibco] in mTeSR1 [StemCell Technologies] and frozen in mFreSR [StemCell Technologies]. Cells were differentiated with 2ng/mL doxycycline [Sigma] at variable conditions as described in (**Supplementary Figure 8**) in either mTeSR1 [StemCell Technologies], RPMI-1640 (microglia) [Gibco] + 10% FBS [Gibco], Williams’ E Medium (hepatocytes) [Gibco] + 10% FBS [Gibco], Immunocult-XF T-Cell Expansion Media (T-cells) [StemCell Technologies], LGM-3 (B-cells) [Lonza], or BrainPhys Media (Astrocytes) [Stem Cell Technologies].

### Flow Cytometry and Cell Sorting

Cells were digested in TrypLE [Gibco] and resuspended in growth media before staining with cell surface markers. The following antibodies were used for analysis and cell sorting: [Microglia: CD11b-FITC, CX3CR1-PE]; [CD8-positive T-Cells: CD3-PerCP-Cy5.5, CD8-FITC]; [T-Regulatory cells: CD3-PerCP-Cy5.5, CD4-PE-Cy7, FOXP3-PE, CD127-V450]; [B-cells: CD19-PE-Cy7, CD27-FITC]; [Hepatocytes: ASGPR1-PE, CD184-APC]; [Astrocytes: CD44-FITC, A2B5-PE]. Cells were sorted and collected on a Sony SH800 FACS for primary screens. For characterization of stable cell lines, cells were stained and analyzed on a BD LSR Fortessa Analyzer flow cytometer. The gating strategy is exemplified in (**Supplementary Figure 6**).

### RNA sequencing

Cells were either collected from FACS (primary screens) or collected directly from culture (refined screens and stable cell line characterization) and were lysed in TRIzol [Invitrogen]. RNA was purified with Direct-zol RNA MicroPrep and RNA MiniPrep kits [Zymo]. Library prep was performed using a SMARTer Stranded Total RNA-Seq Kit v2 - Pico Input Mammalian [TARAKA] (primary screens) and NEBNext Ultra II RNA Kits [NEB] (refined screens and stable cell line characterization). Barcodes were amplified from the prepped cDNA using two alternative primer pairs (**Supplementary Table 5**). Amplicons were sequenced with a MiSeq kit [Illumina] using Illumina TruSeq indexes. Transcriptomes were sequenced on either NextSeq or NovaSeq platforms [Illumina].

### Astrocyte stimulation assays

iAstIIs were differentiated as described in (**Supplementary Figure 8**) and then transferred to imaging dishes for stimulation as previously described [50]. Briefly, glass bottom dishes [Ibidi 81158] were coated in Poly-d-lysine (0.1 mg/mL) for 2 hours at room temperature, washed twice in PBS [Gibco], and coated overnight in fibronectin (10 *µ*g/mL) [Thermo] at 37°C. Differentiated astrocytes were digested in TrypLE [Gibco] for 7-10 minutes, and 40,000-50,000 cells were transferred to coated dishes and maintained for 2 days before stimulation and imaging. Prior to stimulation and imaging the astrocytes were stained with Fluo-4 (1 *µ*g/mL) [Invitrogen] in BrainPhys medium without phenol red [StemCell] and incubated in the dark for at least 25 minutes at 37°C. Cells were then washed with fresh media three times and transferred immediately to a Zeiss Axio 3 Inverted Microscope with CO_2_ (5%) and temperature control (37°C). After staging, basal activity was measured for at least 2 minutes, after which small molecule stimuli were applied.

### Hepatocyte hepatotoxicity assays

iHeps were differentiated as described in (**Supplementary Figure 8**) and then transferred to 96-well plates pre-coated with Matrigel [Corning] and treated with hepatotoxins as previously described [51]. Briefly, after differentiation, 25,000 iHeps, undifferentiated iPSCs, and plateable primary human hepatocytes [ZenBio] were plated in each well and incubated overnight at 37°C. The next day, media was changed to Hepatocyte Medium E (William’s E Medium [Gibco], Maintenance Cocktail B [Gibco], and 0.1*µ*M Dexamethasone [Gibco]) for one day. The following day, media was exchanged and supplemented with hepatotoxins (Acetaminophen at [3.125,6.25,12.5,25,50,100] mM [Spectrum], Nefazodone at [1,3,10,30,100,300] µM [Sigma], and Troglitazone at [1,3,10,30,100,300] *µ*M [Sigma]). Cells were incubated again at 37°C for 24 hours, and viability was measured with CellTiter-Glo Luminescent Cell Viability Assay [Promega].

### Cytotoxic T-cell activation assays

Primary cytotoxic T-cells (Human Peripheral Blood CD4+CD45RA+ T Cells) [StemCell] and iCytoTs were cultured and activated in the same manner. Briefly, cells were incubated in ImmunoCult-XF T Cell Expansion Medium [StemCell] + IL-2 [R&D Systems] with DYNAL Dynabeads Human T-Activator CD3/CD28 for T Cell Expansion and Activation [Gibco] for 3 days. After this incubation, the cells were stained with Celltrace Violet [Invitrogen] and moved into new wells at the concentration of 1M cells/well with fresh media (as above) and grown at 37°C for 11 days, changing media every 2-3 days. Finally, cells were analyzed via flow cytometry. Percent activated was determined by gating cells that had diminished fluorescence after proliferation.

### Regulatory T-cell proliferation suppression assays

iTRegs were co-cultured with activated primary cytotoxic T-cells in variable quantities as previously described [49]. Briefly, iTRegs were differentiated in ImmunoCult-XF T Cell Expansion Medium [StemCell] + IL-2 [R&D Systems] for 4 days and then moved into co-culture with activated and CellTrace Violet [Fisher] stained cytotoxic T-cells and grown at 37°C for 11 days, changing media every 2-3 days. Finally, cells were analyzed via flow cytometry. The percentage of suppression was determined as 100 x [1 - (% of proliferating cells with iTRegs) / (% of proliferating cells without iTRegs)] after applying gates for proliferating v. non-proliferating cells and subtracting auto-fluorescence resulting from unstained iTRegs.

## 5 Acknowledgments

Conceptualization, E.A., J.T., C.G. and G.M.C.; Methodology, E.A. and J.T.; Software, J.T. and E.A.; Formal Analysis J.T.; Investigation, E.A., G.F., S.L.; Writing - Original Draft, E.A. and J.T.; Visualization, E.A. and J.T.; Supervision, C.G. and G.M.C.; Funding Acquisition, E.A and G.M.C. The authors would like to thank Alex Ng, Parastoo Khoshakhlagh, Alexandru Plesa, Christian Kramme, and Jeremy Huang for useful conversations pertaining to this study. This work was funded by the FunGCAT program from the Office of the Director of National Intelligence (ODNI), Intelligence Advanced Research Projects Activity (IARPA), via the Army Research Office (ARO) under Federal Award No. W911NF-17-2-0089 and The EGL Charitable Foundation.

## Notes

### Competing Interest Statement

G.M.C. is an inventor on patents filed by the Presidents and Fellows of Harvard College. Full disclosure for G.M.C. is available at http://arep.med.harvard.edu/gmc/tech.html

### Summary of Updates

Revised manuscript contains more benchmarking comparisons to other tools and some revised claims

## References

[1] Orit Rozenblatt-Rosen, Michael JT Stubbington, Aviv Regev, and Sarah A Teichmann. The human cell atlas: from vision to reality. Nature News, 550(7677):451, 2017.

[2] Kazutoshi Takahashi, Koji Tanabe, Mari Ohnuki, Megumi Narita, Tomoko Ichisaka, Kiichiro Tomoda, and Shinya Yamanaka. Induction of pluripotent stem cells from adult human fibroblasts by defined factors. Cell, 131(5):861–872, 2007.

[3] Jiho Choi, Soohyun Lee, William Mallard, Kendell Clement, Guidantonio Malagoli Tagliazucchi, Hotae Lim, In Young Choi, Francesco Ferrari, Alexander M Tsankov, Ramona Pop, et al. A comparison of genetically matched cell lines reveals the equivalence of human ipscs and escs. Nature biotechnology, 33(11):1173, 2015.

[4] Alex HM Ng, Parastoo Khoshakhlagh, Jesus Eduardo Rojo Arias, Giovanni Pasquini, Kai Wang, Anka Swiersy, Seth L Shipman, Evan Appleton, Kiavash Kiaee, Richie E Kohman, et al. A comprehensive library of human transcription factors for cell fate engineering. Nature Biotechnology, pages 1–10, 2020.

[5] Samantha A Morris, Patrick Cahan, Hu Li, Anna M Zhao, Adrianna K San Roman, Ramesh A Shivdasani, James J Collins, and George Q Daley. Dissecting engineered cell types and enhancing cell fate conversion via cellnet. Cell, 158(4):889–902, 2014.

[6] Xianmei Meng, Amanda Neises, Rui-Jun Su, Kimberly J Payne, Linda Ritter, Daila S Gridley, Jun Wang, Matilda Sheng, KH William Lau, David J Baylink, et al. Efficient reprogramming of human cord blood cd34+ cells into induced pluripotent stem cells with oct4 and sox2 alone. Molecular Therapy, 20(2):408–416, 2012.

[7] Jonathan Sagal, Xiping Zhan, Jinchong Xu, Jessica Tilghman, Senthilkumar S Karuppagounder, Li Chen, Valina L Dawson, Ted M Dawson, John Laterra, and Mingyao Ying. Proneural transcription factor atoh1 drives highly efficient differentiation of human pluripotent stem cells into dopaminergic neurons. Stem cells translational medicine, 3(8):888–898, 2014.

[8] Volker Busskamp, Nathan E Lewis, Patrick Guye, Alex HM Ng, Seth L Shipman, Susan M Byrne, Neville E Sanjana, Jernej Murn, Yinqing Li, Shangzhong Li, et al. Rapid neurogenesis through transcriptional activation in human stem cells. Molecular systems biology, 10(11), 2014.

[9] Sven Heinz, Christopher Benner, Nathanael Spann, Eric Bertolino, Yin C Lin, Peter Laslo, Jason X Cheng, Cornelis Murre, Harinder Singh, and Christopher K Glass. Simple combinations of lineage-determining transcription factors prime cis-regulatory elements required for macrophage and b cell identities. Molecular cell, 38(4):576–589, 2010.

[10] Madeline A Lancaster and Juergen A Knoblich. Organogenesis in a dish: modeling development and disease using organoid technologies. Science, 345(6194):1247125, 2014.

[11] Marius Ader and Elly M Tanaka. Modeling human development in 3d culture. Current opinion in cell biology, 31:23–28, 2014.

[12] Priyalakshmi Viswanathan, Matthew G Ondeck, Somyot Chirasatitsin, Kamolchanok Ngamkham, Gwendolen C Reilly, Adam J Engler, and Giuseppe Battaglia. 3d surface topology guides stem cell adhesion and differentiation. Biomaterials, 52:140–147, 2015.

[13] Thomas Graf and Tariq Enver. Forcing cells to change lineages. Nature, 462(7273):587, 2009.

[14] Thomas Vierbuchen, Austin Ostermeier, Zhiping P Pang, Yuko Kokubu, Thomas C Südhof, and Marius Wernig. Direct conversion of fibroblasts to functional neurons by defined factors. Nature, 463(7284):1035, 2010.

[15] Masaki Ieda, Ji-Dong Fu, Paul Delgado-Olguin, Vasanth Vedantham, Yohei Hayashi, Benoit G Bruneau, and Deepak Srivastava. Direct reprogramming of fibroblasts into functional cardiomyocytes by defined factors. Cell, 142(3):375–386, 2010.

[16] Pengyu Huang, Zhiying He, Shuyi Ji, Huawang Sun, Dao Xiang, Changcheng Liu, Yiping Hu, Xin Wang, and Lijian Hui. Induction of functional hepatocyte-like cells from mouse fibroblasts by defined factors. Nature, 475(7356):386, 2011.

[17] Owen JL Rackham, Jaber Firas, Hai Fang, Matt E Oates, Melissa L Holmes, Anja S Knaupp, Harukazu Suzuki, Christian M Nefzger, Carsten O Daub, Jay W Shin, et al. A predictive computational framework for direct reprogramming between human cell types. Nature genetics, 48(3):331, 2016.

[18] András Hartmann, Satoshi Okawa, Gaia Zaffaroni, and Antonio del Sol. Seesawpred: A web application for predicting cell-fate determinants in cell differentiation. Scientific reports, 8(1):13355, 2018.

[19] Satoshi Okawa, Carmen Saltó, Srikanth Ravichandran, Shanzheng Yang, Enrique M Toledo, Ernest Arenas, and Antonio del Sol. Transcriptional synergy as an emergent property defining cell subpopulation identity enables population shift. Nature communications, 9(1):2595, 2018.

[20] Christian Kramme, Alexandru Plesa, Helen H Wang, Bennett Wolf, Merrick Smela, Xiaoge Guo, Richie Kohman, Pranam Chatterjee, and George Church. Stampscreen: An integrated pipeline for mammalian genetic screening. Available at SSRN 3875773.

[21] Patrick Cahan, Hu Li, Samantha A Morris, Edroaldo Lummertz Da Rocha, George Q Daley, and James J Collins. Cellnet: network biology applied to stem cell engineering. Cell, 158(4):903–915, 2014.

[22] Sascha Jung, Evan Appleton, Muhammad Ali, George M Church, and Antonio Del Sol. A computer-guided design tool to increase the efficiency of cellular conversions. Nature communications, 12(1):1–12, 2021.

[23] Ning Leng, John A Dawson, James A Thomson, Victor Ruotti, Anna I Rissman, Bart MG Smits, Jill D Haag, Michael N Gould, Ron M Stewart, and Christina Kendziorski. Ebseq: an empirical bayes hierarchical model for inference in rna-seq experiments. Bioinformatics, 29(8):1035–1043, 2013.

[24] Nurcan Tuncbag, Sara JC Gosline, Amanda Kedaigle, Anthony R Soltis, Anthony Gitter, and Ernest Fraenkel. Network-based interpretation of diverse high-throughput datasets through the omics integrator software package. PLoS computational biology, 12(4):e1004879, 2016.

[25] Timothy L Bailey. Dreme: motif discovery in transcription factor chip-seq data. Bioinformatics, 27(12):1653–1659, 2011.

[26] Yuchun Guo, Kevin Tian, Haoyang Zeng, Xiaoyun Guo, and David Kenneth Gifford. A novel k-mer set memory (ksm) motif representation improves regulatory variant prediction. Genome research, 28(6):891–900, 2018.

[27] Philip Machanick and Timothy L Bailey. Meme-chip: motif analysis of large dna datasets. Bioinformatics, 27(12):1696–1697, 2011.

[28] Jennifer Hammelman and David K Gifford. Discovering differential genome sequence activity with interpretable and efficient deep learning. PLoS Computational Biology, 17(8):e1009282, 2021.

[29] Ivan Berest, Christian Arnold, Armando Reyes-Palomares, Giovanni Palla, Kasper Dindler Rasmussen, Holly Giles, Peter-Martin Bruch, Wolfgang Huber, Sascha Dietrich, Kristian Helin, et al. Quantification of differential transcription factor activity and multiomics-based classification into activators and repressors: difftf. Cell reports, 29(10):3147–3159, 2019.

[30] Quan Xu, Georgios Georgiou, Siebren Frölich, Maarten van der Sande, Gert Jan C Veenstra, Huiqing Zhou, and Simon J van Heeringen. Ananse: an enhancer network-based computational approach for predicting key transcription factors in cell fate determination. Nucleic acids research, 49(14):7966–7985, 2021.

[31] Florian Schmidt, Nina Gasparoni, Gilles Gasparoni, Kathrin Gianmoena, Cristina Cadenas, Julia K Polansky, Peter Ebert, Karl Nordström, Matthias Barann, Anupam Sinha, et al. Combining transcription factor binding affinities with open-chromatin data for accurate gene expression prediction. Nucleic acids research, 45(1):54–66, 2017.

[32] David T Jones. Setting the standards for machine learning in biology. Nature Reviews Molecular Cell Biology, 20(11):659–660, 2019.

[33] Kevin K Yang, Zachary Wu, and Frances H Arnold. Machine-learning-guided directed evolution for protein engineering. Nature methods, 16(8):687–694, 2019.

[34] Gökcen Eraslan, Žiga Avsec, Julien Gagneur, and Fabian J Theis. Deep learning: new computational modelling techniques for genomics. Nature Reviews Genetics, 20(7):389–403, 2019.

[35] Pierce J Ogden, Eric D Kelsic, Sam Sinai, and George M Church. Comprehensive aav capsid fitness landscape reveals a viral gene and enables machine-guided design. Science, 366(6469):1139–1143, 2019.

[36] Aderonke F Ajayi-Smith, Pauline J van der Watt, and Virna D Leaner. Interfering with nuclear transport as a means of interrupting transcription factor activity in cancer. Critical Reviews™ in Eukaryotic Gene Expression, 29(5), 2019.

[37] Yaser Atlasi and Hendrik G. Stunnenberg. The interplay of epigenetic marks during stem cell differentiation and development. Nature Reviews Genetics, 18(11):643–658, Nov 2017.

[38] Ryan Raisner, Samir Kharbanda, Lingyan Jin, Edwin Jeng, Emily Chan, Mark Merchant, Peter M. Haverty, Russell Bainer, Tommy Cheung, David Arnott, E. Megan Flynn, F. Anthony Romero, Steven Magnuson, and Karen E. Gascoigne. Enhancer activity requires cbp/p300 bromodomain-dependent histone h3k27 acetylation. Cell Reports, 24(7):1722–1729, 2018.

[39] Bing Ren, François Robert, John J. Wyrick, Oscar Aparicio, Ezra G. Jennings, Itamar Simon, Julia Zeitlinger, Jörg Schreiber, Nancy Hannett, Elenita Kanin, Thomas L. Volkert, Christopher J. Wilson, Stephen P. Bell, and Richard A. Young. Genome-wide location and function of dna binding proteins. Science, 290(5500):2306–2309, 2000.

[40] ENCODE Project Consortium et al. The encode (encyclopedia of dna elements) project. Science, 306(5696):636– 640, 2004.

[41] Ron Edgar, Michael Domrachev, and Alex E Lash. Gene expression omnibus: Ncbi gene expression and hybridization array data repository. Nucleic acids research, 30(1):207–210, 2002.

[42] Aziz Khan, Oriol Fornes, Arnaud Stigliani, Marius Gheorghe, Jaime A Castro-Mondragon, Robin Van Der Lee, Adrien Bessy, Jeanne Cheneby, Shubhada R Kulkarni, Ge Tan, et al. Jaspar 2018: update of the open-access database of transcription factor binding profiles and its web framework. Nucleic acids research, 46(D1):D260–D266, 2018.

[43] Aziz Khan, Oriol Fornes, Arnaud Stigliani, Marius Gheorghe, Jaime A Castro-Mondragon, Robin van der Lee, Adrien Bessy, Jeanne Cheneby, Shubhada R Kulkarni, Ge Tan, et al. Jaspar 2018: update of the open-access database of transcription factor binding profiles and its web framework. Nucleic acids research, 46(D1):D260–D266, 2017.

[44] Constantinos Chronis, Petko Fiziev, Bernadett Papp, Stefan Butz, Giancarlo Bonora, Shan Sabri, Jason Ernst, and Kathrin Plath. Cooperative binding of transcription factors orchestrates reprogramming. Cell, 168(3):442–459.e20, 2017.

[45] Gregory J Fonseca, Jenhan Tao, Emma M Westin, Sascha H Duttke, Nathanael J Spann, Tobias Strid, Zeyang Shen, Joshua D Stender, Mashito Sakai, Verena M Link, et al. Diverse motif ensembles specify non-redundant dna binding activities of ap-1 family members in macrophages. Nature communications, 10(1):414, 2019.

[46] Samuel A Lambert, Arttu Jolma, Laura F Campitelli, Pratyush K Das, Yimeng Yin, Mihai Albu, Xiaoting Chen, Jussi Taipale, Timothy R Hughes, and Matthew T Weirauch. The human transcription factors. Cell, 172(4):650–665, 2018.

[47] Bárbara González, Sabine Denzel, Brigitte Mack, Marcus Conrad, and Olivier Gires. Epcam is involved in maintenance of the murine embryonic stem cell phenotype. Stem cells, 27(8):1782–1791, 2009.

[48] Yingyao Zhou, Bin Zhou, Lars Pache, Max Chang, Alireza Hadj Khodabakhshi, Olga Tanaseichuk, Christopher Benner, and Sumit K Chanda. Metascape provides a biologist-oriented resource for the analysis of systems-level datasets. Nature communications, 10(1):1–10, 2019.

[49] Yehezqel Elyahu, Idan Hekselman, Inbal Eizenberg-Magar, Omer Berner, Itai Strominger, Maya Schiller, Kritika Mittal, Anna Nemirovsky, Ekaterina Eremenko, Assaf Vital, et al. Aging promotes reorganization of the cd4 t cell landscape toward extreme regulatory and effector phenotypes. Science advances, 5(8):eaaw8330, 2019.

[50] Marita Grønning Hansen, Daniel Tornero, Isaac Canals, Henrik Ahlenius, and Zaal Kokaia. In vitro functional characterization of human neurons and astrocytes using calcium imaging and electrophysiology. In Neural Stem Cells, pages 73–88. Springer, 2019.

[51] Jingtao Lu, Shannon Einhorn, Lata Venkatarangan, Manda Miller, David A Mann, Paul B Watkins, and Edward LeCluyse. Morphological and functional characterization and assessment of ipsc-derived hepatocytes for in vitro toxicity testing. Toxicological Sciences, 147(1):39–54, 2015.

[52] Kazuo Takayama, Yuta Morisaki, Shuichi Kuno, Yasuhito Nagamoto, Kazuo Harada, Norihisa Furukawa, Manami Ohtaka, Ken Nishimura, Kazuo Imagawa, Fuminori Sakurai, et al. Prediction of interindividual differences in hepatic functions and drug sensitivity by using human ips-derived hepatocytes. Proceedings of the National Academy of Sciences, 111(47):16772–16777, 2014.

[53] Thomas Barthlott, Halima Moncrieffe, Marc Veldhoen, Christopher J Atkins, Jillian Christensen, Anne O’Garra, and Brigitta Stockinger. Cd25+ cd4+ t cells compete with naive cd4+ t cells for il-2 and exploit it for the induction of il-10 production. International immunology, 17(3):279–288, 2005.

[54] Ben Langmead and Steven L Salzberg. Fast gapped-read alignment with Bowtie 2. Nat Methods, 9(4):357–359, 2012.

[55] Peter Sonneveld and Martin B Van Gijzen. Idr (s): A family of simple and fast algorithms for solving large nonsymmetric systems of linear equations. SIAM Journal on Scientific Computing, 31(2):1035–1062, 2009.

[56] Robert Tibshirani. Regression Shrinkage and Selection Via the Lasso. Journal of the Royal Statistical Society: Series B (Methodological), 58(1):267–288, jan 1996.

[57] Wes McKinney et al. pandas: a foundational python library for data analysis and statistics. Python for high performance and scientific computing, 14(9):1–9, 2011.

[58] Travis E Oliphant. A guide to NumPy, volume 1. Trelgol Publishing USA, 2006.

[59] Pauli Virtanen, Ralf Gommers, Travis E. Oliphant, Matt Haberland, Tyler Reddy, David Cournapeau, Evgeni Burovski, Pearu Peterson, Warren Weckesser, Jonathan Bright, Stéfan J. van der Walt, Matthew Brett, Joshua Wilson, K. Jarrod Millman, Nikolay Mayorov, Andrew R. J. Nelson, Eric Jones, Robert Kern, Eric Larson, C J Carey, İlhan Polat, Yu Feng, Eric W. Moore, Jake VanderPlas, Denis Laxalde, Josef Perktold, Robert Cimrman, Ian Henriksen, E. A. Quintero, Charles R. Harris, Anne M. Archibald, Antônio H. Ribeiro, Fabian Pedregosa, Paul van Mulbregt, and SciPy 1.0 Contributors. SciPy 1.0: Fundamental Algorithms for Scientific Computing in Python. Nature Methods, 17:261–272, 2020.

[60] F. Pedregosa, G. Varoquaux, A. Gramfort, V. Michel, B. Thirion, O. Grisel, M. Blondel, P. Prettenhofer, R. Weiss, V. Dubourg, J. Vanderplas, A. Passos, D. Cournapeau, M. Brucher, M. Perrot, and E. Duchesnay. Scikit-learn: Machine learning in Python. Journal of Machine Learning Research, 12:2825–2830, 2011.

[61] Peter J.A. Cock, Tiago Antao, Jeffrey T. Chang, Brad A. Chapman, Cymon J. Cox, Andrew Dalke, Iddo Friedberg, Thomas Hamelryck, Frank Kauff, Bartek Wilczynski, and Michiel J.L. De Hoon. Biopython: Freely available Python tools for computational molecular biology and bioinformatics. Bioinformatics, 25(11):1422–1423, 2009.

[62] Wolfgang Huber, Vincent J Carey, Robert Gentleman, Simon Anders, Marc Carlson, Benilton S Carvalho, Hector Corrada Bravo, Sean Davis, Laurent Gatto, Thomas Girke, et al. Orchestrating high-throughput genomic analysis with bioconductor. Nature methods, 12(2):115–121, 2015.

[63] Michael I. Love, Wolfgang Huber, and Simon Anders. Moderated estimation of fold change and dispersion for RNA-seq data with DESeq2. Genome Biology, 15(12):1–21, 2014.

