## Supplementary Information for "Machine-guided cell-fate engineering"

### MACHINE-GUIDED CELL-FATE ENGINEERING
### SUPPLEMENTARY INFORMATION

Evan Appleton[1,2,*], Jenhan Tao[3,*], Greg Fonseca[3], Songlei Liu[1,2], Christopher Glass[3], and George Church[1,2]
[1]Wyss Institute for Biologically Inspired Engineering at Harvard University, Boston, Massachusetts, USA
[2]Department of Genetics, Harvard Medical School, Boston, Massachusetts, USA
[3]University of California, San Diego, San Diego, CA
[*]These authors contributed equally

### 1 Flow cytometry analysis

Gating raw data to determine positive and negative populations for flow cyomtery analysis is a critical and necessary step of the *CellCartographer* pipeline. The gating strategy is tweaked for three different modes of screens/characterization: primary screens, polyclonal cell lines, and monoclonal cell lines. These considerations are exemplified in **Supplementary Figure 6** using ggplot2 [1].

For primary screens, gating was done to remove electonric noise and separate singlets from doublets (**Supplementary Figure 6d**), and then gates were drawn around unstained and stained iPSC controls. For refined cell lines that were built with fewer TFs, the same analysis was performed because these populations were still of mixed genetic identity and therefore cell size and shape might still be variable. For clonal cell populations, however, cell size and shape was nearly unanimous. Therefore, as a final step to clean up outliers, we applied a k-means filter on forward and side scatter after filtering out singlet, doublets, and electronic noise (**Supplementary Figure 6a,b**).

Twelve total cell types were screened in the primary screening process. This was done to cover a diverse spread of cell types of potential diagnostic or therapeutic relevance from all germ layers. Furthermore we wanted to demonstrate that *CellCartographer* can generate strong results for cell sub-types (i.e. T-cell subtypes and epithelial cell types (**Supplementary Figure 7**).

We found that for all of the cell types we assayed, there were small populations that were clearly different from the stem cell controls. For nine of the cell types (type II astrocytes, B-cells, CD8+ T-cells, microglia, hepatocytes, regulatory T-cells, osteoblasts, delta-gamma T-cells, and macrophages) we found small double-positive populations when assayed in mTeSR as described. For the remaining three cell types (bronchial epithelial cells, kidney epithelial cells, and mammary epithelial cells), we clearly saw changes in key markers compared to stem cells, but there was some difficulty in determining exactly how to interpret these differences. Namely, our stem cells appeared to share some markers with epithelial cells (**Supplementary Figure 6d-f**). For example, it seemed that CD24 was highly expressed in our stem cells, but seemed to lose this marker dramatically when TFs for kindey and mammary epithelial cells were expressed. This also seemed to be the case to a lesser to degree with CD45 for bronchial epithelial cells. These cases show clear differences between controls and the experiments, but the differences are hard to interpret. This challenge did not have an apparently obvious solution because our ability to identify clear cell-surface markers across these types from the literature was not fruitful. Ergo, one limitation of the *CellCatorgrapher* pipeline is the ability to identify unambiguous surface marker anitbodies for cell types of interest - if well-established canonical markers for a given cell type cannot be found, the *CellCartographer* pipeline will require extra workarounds, such as sequencing more cell populations or using single-cell sequencing.

#### 2 Cell line differentiation

Differentiation conditions for primary screens (**Figure 1**) and refined cell lines (**Supplementary Figure 8a**) was first kept as simple as possible as a benchmark of success of the TFs to perform differentiation operations. However, we wanted to explore if changing the differentiation medium to the ideal medium for the final cell type was successful in accelerating the differentiation process along-side the TFs. To determine this, we identified media that were suitable for maintaining and expanding primary cells of the destination cell type and performed differentiations with DOX in these medias one day after plating stem cells (**Supplementary Figure 8b-f**). Furthermore, since we wanted to avoid introducing the variable of passaging in measuring the differentiation conditions, we had to optimize the number of cells that were initially plated and how many days they would expand. Given that this was variable for each cell type, we generally seeded at a low quantity of cell (50,000 cells) given that they were to grow in these containers for 6+ days.

A special consideration was given to lymphocytes (T-cells and B-cells) — given that these types of cells often grow best in at higher density and different oxygen concentrations [2], we optimized the seeding density for reproducibility for these cell types and found that the most consistent growth occurred at higher initial seeding density (250,000 cells). This did not cause issues for confluency because as these cell differentiate, they will naturally start to dissociate from the plate surface as they are undergoing a change from surface-adherent, to non surface-adherent cell types. This differentiation culturing methodology could probably benefit from even more cell culture method optimization.

Differentiation of stable differentiation-inducible cell lines in mTeSR even for stable refined cell lines showed dramatic improvement of differentiation efficiency compared to primary screens (**Supplementary Figure 9**) after 6 days of differentiation. However, even for high-performance clones, this percentage was still often in the neighborhood of 20-25% double-positive after 6 days differentiated in mTeSR.

However, when we consider that these cells are still somewhat naive and not fully developed after a short time in stem cell medium, we looked at the percentage of cells that were positive for either cell surface marker compared to those that were clearly double-negative (**Supplementary Figure 10**). When we look at the differentiation efficiencies in this way, most high-performing clonal lines were in the neighborhood of 50% differentiated (**Supplementary Figure 12**).

When we then examine the differentiation efficiency of our cell lines in media suitable for culturing these individual cell types, in most cases we that differentiation into cells positive for at least one cell-surface marker is often much closer to 90-100% after only 6 days of differentiation.

### References


[1] Hadley Wickham. *ggplot2: Elegant Graphics for Data Analysis*. Springer-Verlag New York, 2016.

[2] Kondala R Atkuri, Leonard A Herzenberg, Anna-Kaisa Niemi, Tina Cowan, and Leonore A Herzenberg. Importance of culturing primary lymphocytes at physiological oxygen levels. *Proceedings of the National Academy of Sciences*, 104(11):4547–4552, 2007.

[3] Patrick Cahan, Hu Li, Samantha A Morris, Edroaldo Lummertz Da Rocha, George Q Daley, and James J Collins. Cellnet: network biology applied to stem cell engineering. *Cell*, 158(4):903–915, 2014.

[4] Owen JL Rackham, Jaber Firas, Hai Fang, Matt E Oates, Melissa L Holmes, Anja S Knaupp, Harukazu Suzuki, Christian M Nefzger, Carsten O Daub, Jay W Shin, et al. A predictive computational framework for direct reprogramming between human cell types. *Nature genetics*, 48(3):331, 2016.

[5] Sascha Jung, Evan Appleton, Muhammad Ali, George M Church, and Antonio Del Sol. A computer-guided design tool to increase the efficiency of cellular conversions. *Nature communications*, 12(1):1–12, 2021.

[6] Ning Leng, John A Dawson, James A Thomson, Victor Ruotti, Anna I Rissman, Bart MG Smits, Jill D Haag, Michael N Gould, Ron M Stewart, and Christina Kendziorski. Ebseq: an empirical bayes hierarchical model for inference in rna-seq experiments. *Bioinformatics*, 29(8):1035–1043, 2013.

[7] Nurcan Tuncbag, Sara JC Gosline, Amanda Kedaigle, Anthony R Soltis, Anthony Gitter, and Ernest Fraenkel. Network-based interpretation of diverse high-throughput datasets through the omics integrator software package. *PLoS computational biology*, 12(4):e1004879, 2016.

[8] Sven Heinz, Christopher Benner, Nathanael Spann, Eric Bertolino, Yin C Lin, Peter Laslo, Jason X Cheng, Cornelis Murre, Harinder Singh, and Christopher K Glass. Simple combinations of lineage-determining transcription factors prime cis-regulatory elements required for macrophage and b cell identities. *Molecular cell*, 38(4):576–589, 2010.

[9] Timothy L Bailey. Dreme: motif discovery in transcription factor chip-seq data. *Bioinformatics*, 27(12):1653–1659, 2011.

[10] Yuchun Guo, Kevin Tian, Haoyang Zeng, Xiaoyun Guo, and David Kenneth Gifford. A novel k-mer set memory (ksm) motif representation improves regulatory variant prediction. *Genome research*, 28(6):891–900, 2018.

[11] Philip Machanick and Timothy L Bailey. Meme-chip: motif analysis of large dna datasets. *Bioinformatics*, 27(12):1696–1697, 2011.

[12] Jennifer Hammelman and David K Gifford. Discovering differential genome sequence activity with interpretable and efficient deep learning. *PLoS Computational Biology*, 17(8):e1009282, 2021.

[13] Ivan Berest, Christian Arnold, Armando Reyes-Palomares, Giovanni Palla, Kasper Dindler Rasmussen, Holly Giles, Peter-Martin Bruch, Wolfgang Huber, Sascha Dietrich, Kristian Helin, et al. Quantification of differential transcription factor activity and multiomics-based classification into activators and repressors: diffTF. *Cell reports*, 29(10):3147–3159, 2019.

[14] Quan Xu, Georgios Georgiou, Siebren Frölich, Maarten van der Sande, Gert Jan C Veenstra, Huiqing Zhou, and Simon J van Heeringen. Ananse: an enhancer network-based computational approach for predicting key transcription factors in cell fate determination. *Nucleic acids research*, 49(14):7966–7985, 2021.

[15] Florian Schmidt, Nina Gasparoni, Gilles Gasparoni, Kathrin Gianmoena, Cristina Cadenas, Julia K Polansky, Peter Ebert, Karl Nordström, Matthias Barann, Anupam Sinha, et al. Combining transcription factor binding affinities with open-chromatin data for accurate gene expression prediction. *Nucleic acids research*, 45(1):54–66, 2017.

[16] Jennifer Hammelman, Tulsi Patel, Michael Closser, Hynek Wichterle, and David Gifford. Ranking reprogramming factors for cell differentiation. *Nature Methods*, 19(7):812–822, 2022.

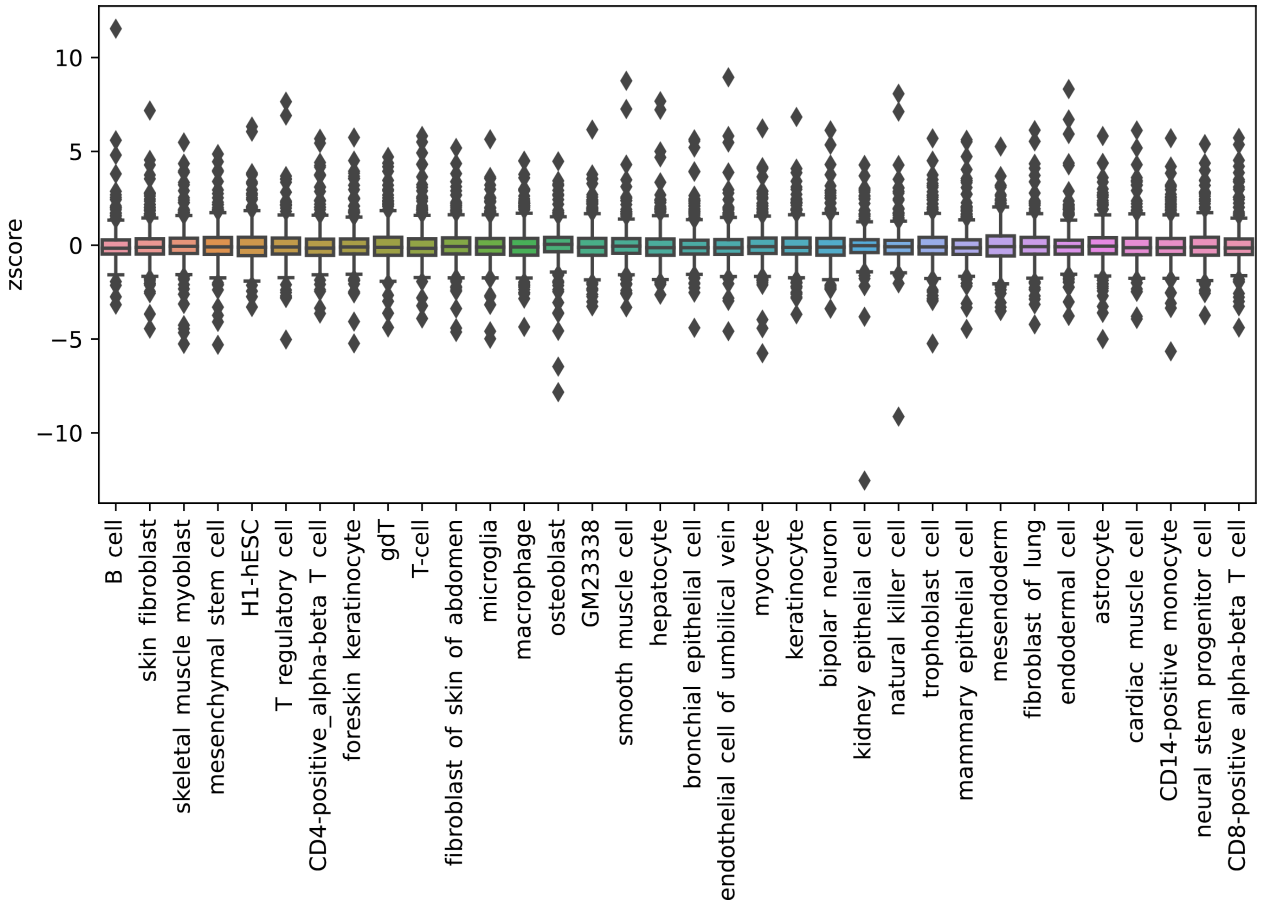


**Supplementary Figure 1: Identification of constitutively active TFs.** Distribution of model coefficients after z-score normalization for each cell type

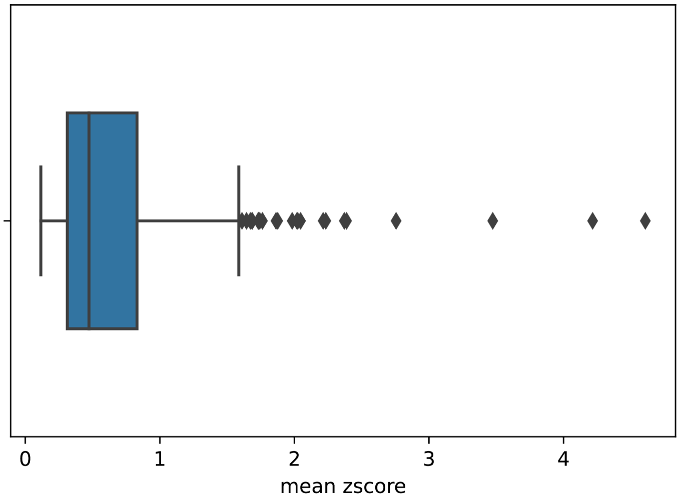


**Supplementary Figure 2: Distribution of mean absolute z-score for each motif across all cell types and tissues.**

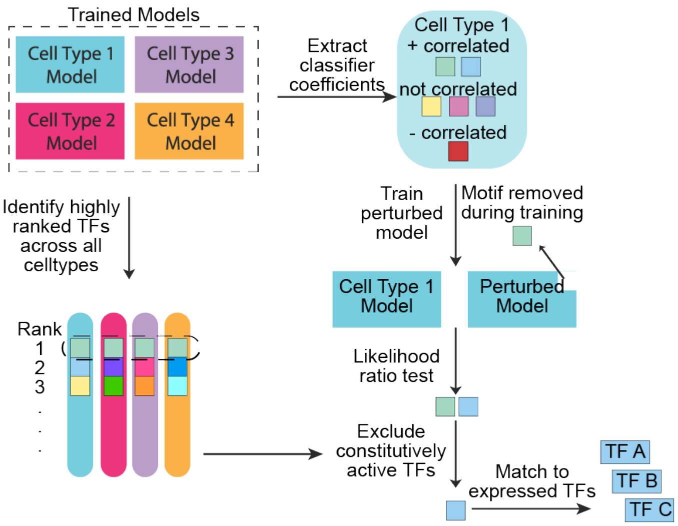


**Supplementary Figure 3:** Algorithmic workflow diagram for CellCartographer.

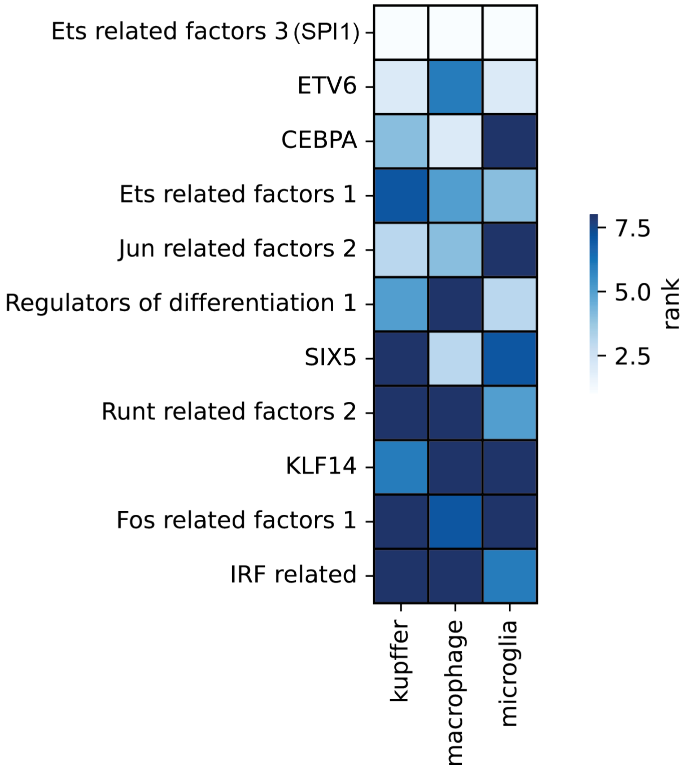


**Supplementary Figure 4:** Comparison of top-8 results of closely related cells with closely overlapping transcriptomes - marcrophage, microglia, and kuppfer cells.

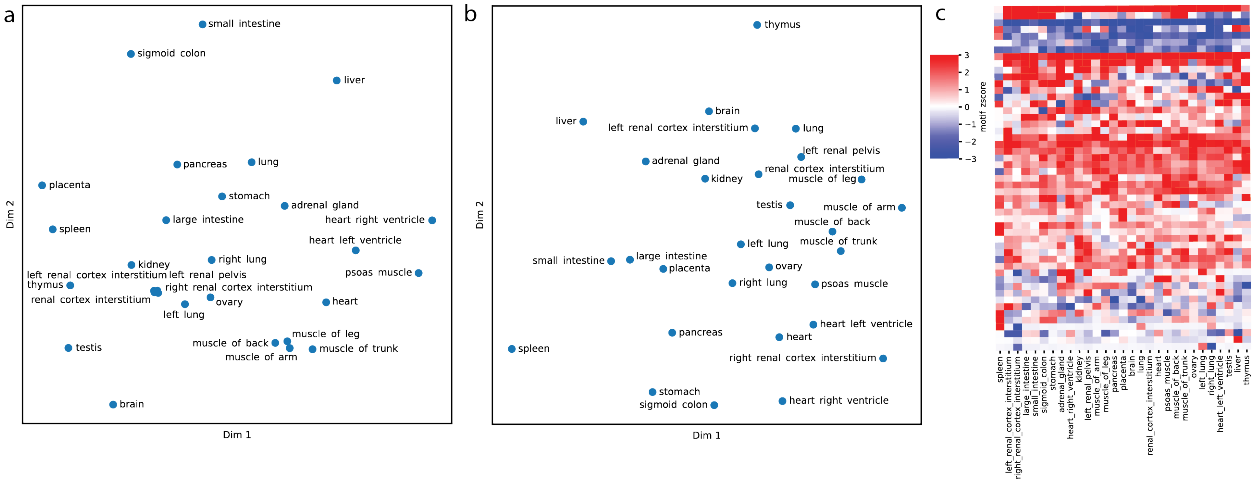


**Supplementary Figure 5: Computational analysis of 29 tissue types with *CellCartographer*.a.** Multidimensional scaling of the similarity in gene expression between different cell types. **b.** Multidimensional scaling of the similarity in TFs correlated with open chromatin. **c.** Motifs correlated (red) and anti-correlated (blue) with open chromatin vary across 29 tissue types analyzed.

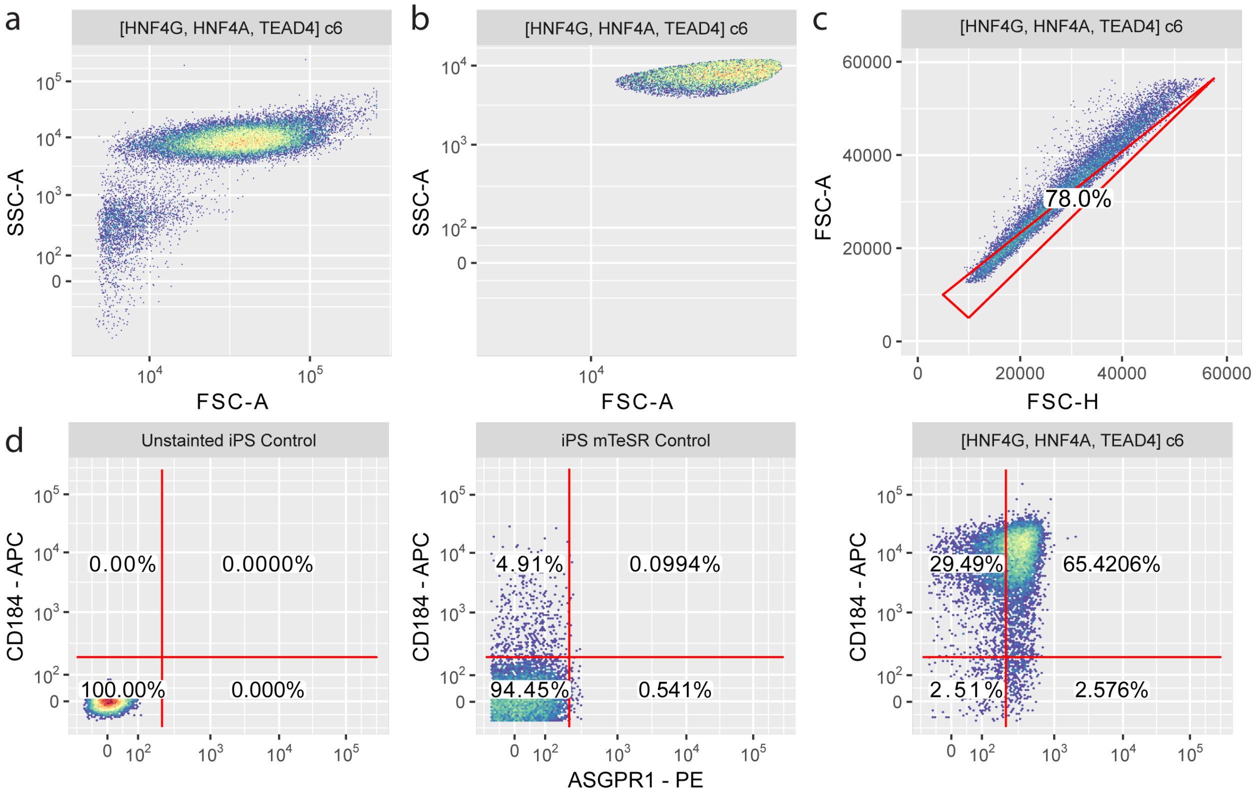


**Supplementary Figure 6: Cytometry gating strategy exemplified on iHep cell line. a.** Initial cell populations included all events on FSC and SSC. **b.** Cellular size/shape outlier events were filtered using a k-means filter for FSC and SSC. **c.** Singlet cells were identified with a linear filter on FSC-A and FSC-H. **d.** Unstained iPSCs. **e.** Gates for double-positive events drawn around iPSCs stained with cell-surface markers. **f.** Same gates for double-positive events drawn around iHeps to determine percent double-positive events.

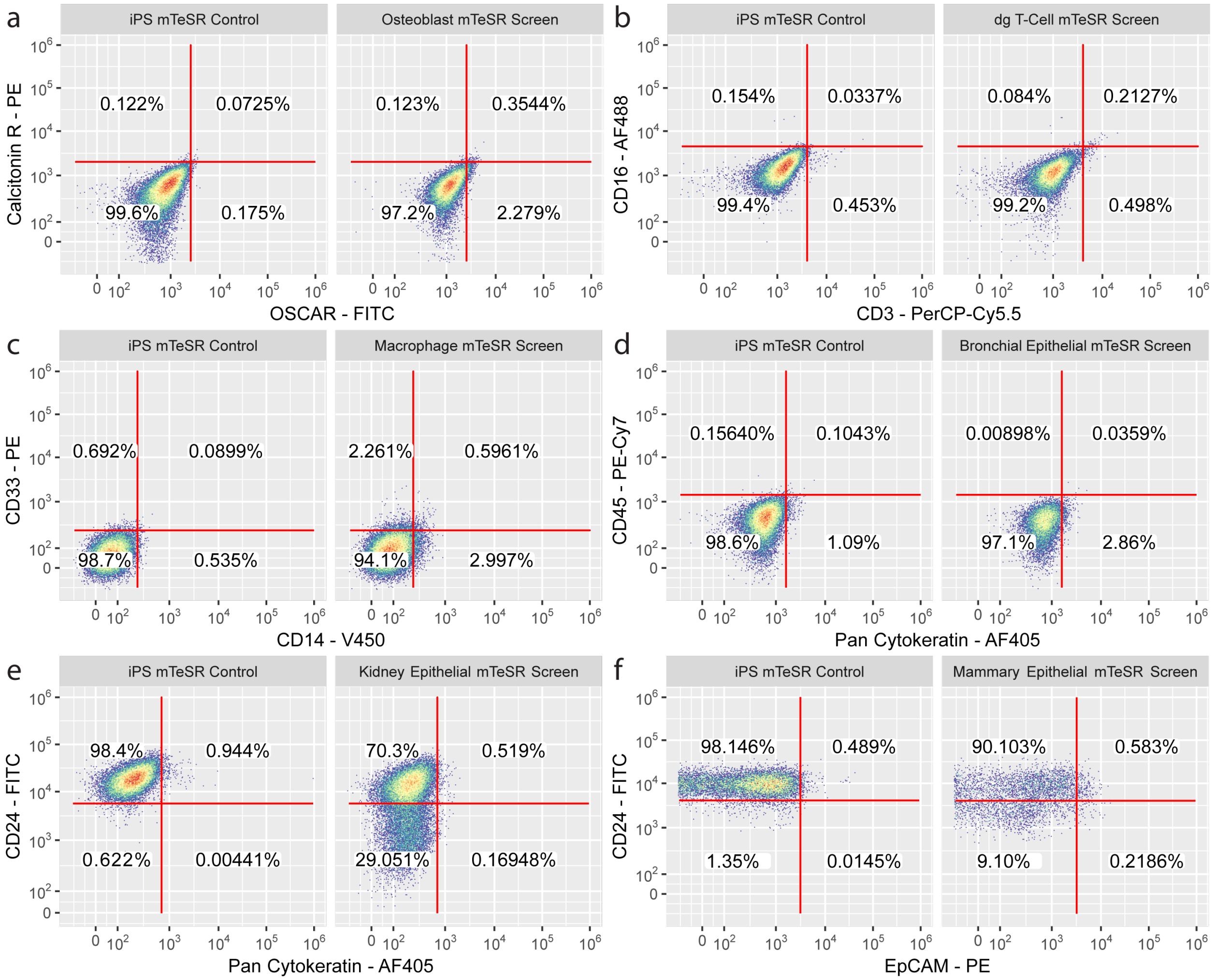


**Supplementary Figure 7: Primary pooled screens for additional 6 cell types.** For each cell type, a negative antibody stain for iPSCs without TFs (LEFT) and the cell population with induced TFs (RIGHT) is shown. **a.** Osteoblasts (mesoderm) **b.** delta-gamma T-cells (mesoderm) **c.** Macrophages (mesoderm) **d.** Bronchial epithelial cells (mesoderm). **e.** Kidney epithelial cells (mesoderm) **f.** Mammary epithelial cells (mesoderm).

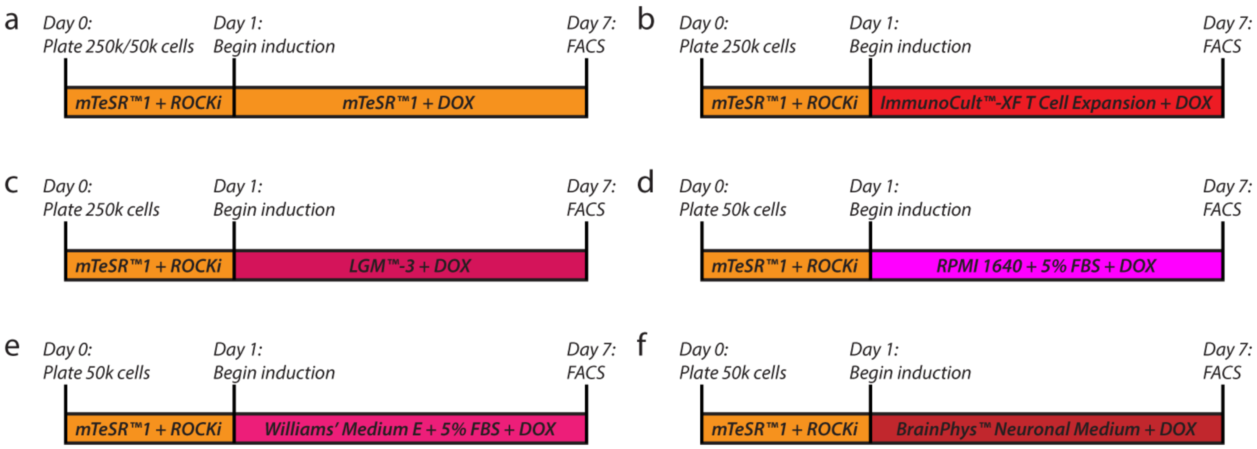


**Supplementary Figure 8: Stable cell line differentiation conditions.** Stem cell lines differentiated into adherent cell types were seeded with 50k cells and stem cell lines differentiated into non-adherent (i.e. lymphocyte types) were seeded with 250k cells, each for 6 days prior to flow cytometry. **a.** All stable cell lines were tested by differentiating cells in mTeSR. **b.** T-cells were differentiated in ImmunoCult -XF T Cell Expansion medium. **c.** B-cells were differentiated in LGM-3 medium. **d.** Microglia were differentiated in RPMI medium + 5% FBS. **e.** Hepatocytes were differentiated in Williams' Medium E + 5% FBS. **f.** Type II astrocytes were differentiated in BrainPhys Neuronal Medium.

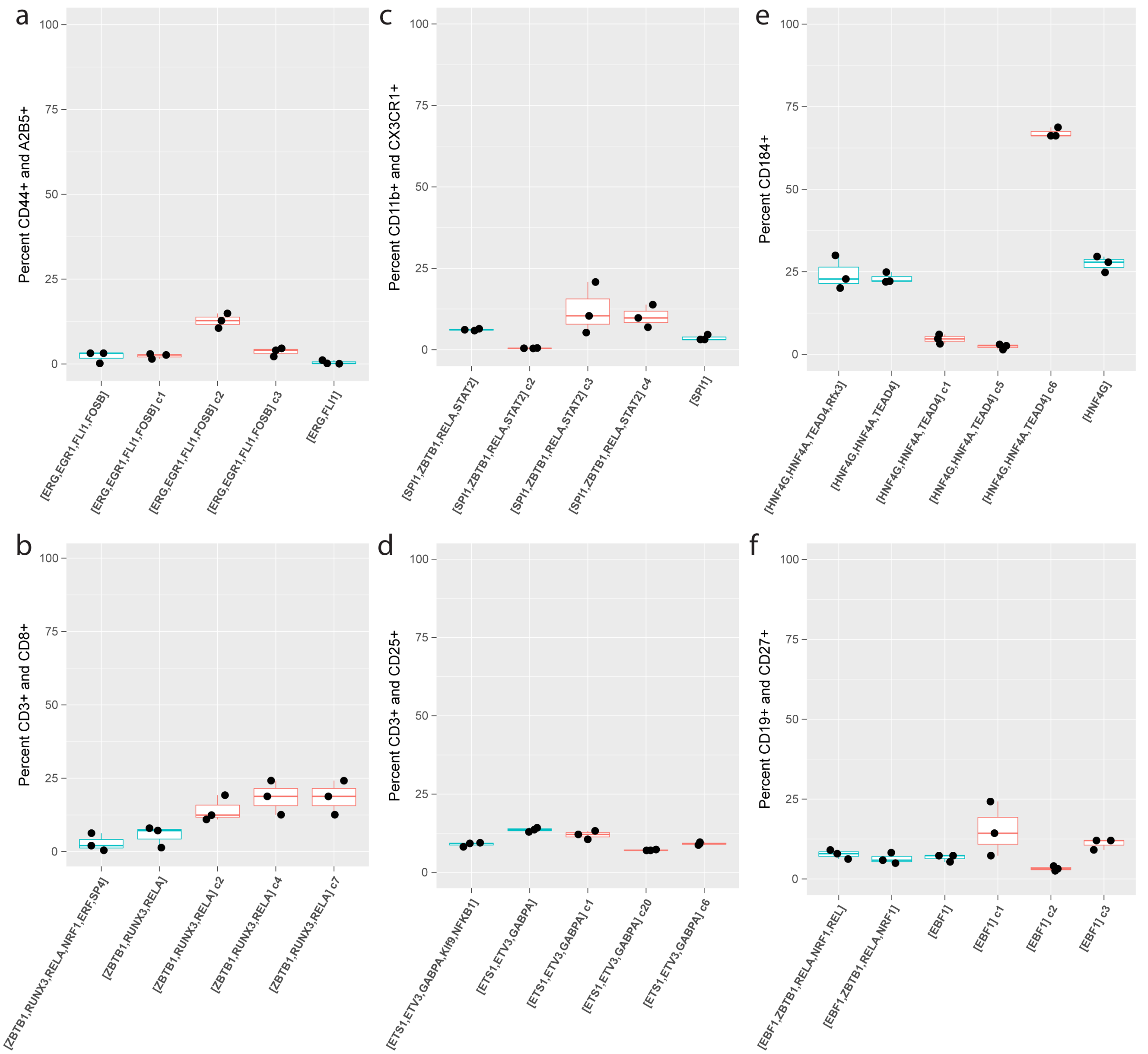


**Supplementary Figure 9: Cell line differentiation for double-positive cell surface markers after 6 days of differentiation in mTeSR + DOX.** For each cell type, we show percent double-positive for FACS analysis of canonical markers for non-clonal (turquoise) and mono-clonal (red) cell lines. **a.** Type II Astrocytes **b.** CD8-positive T-cells **c.** Microglia **d.** Regulatory T-cells **e.** Hepatocytes and **f.** B-cells (mesoderm).

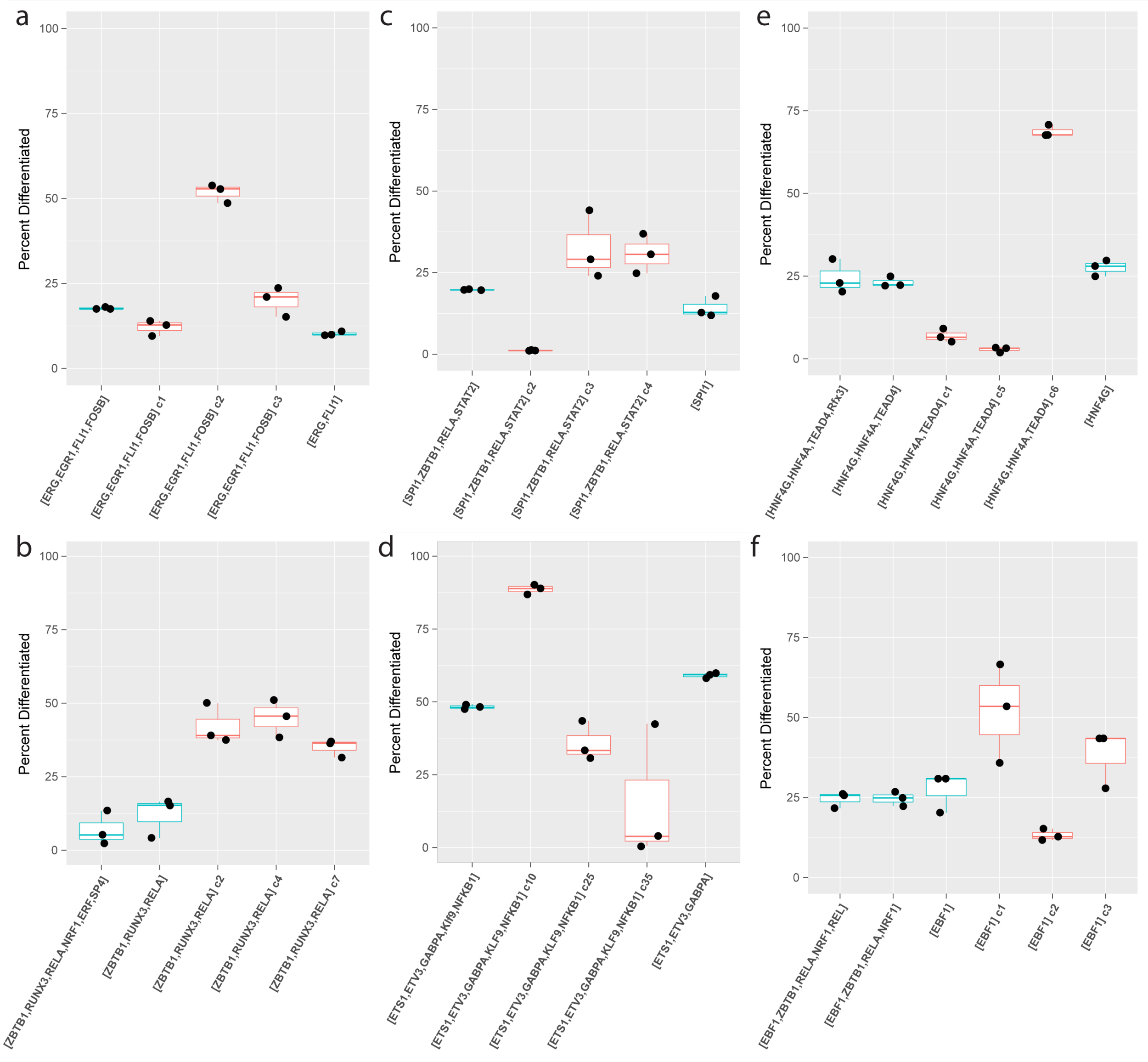


**Supplementary Figure 10: Cell line differentiation for either one or both cell surface markers in cell-type-specific growth medium after 6 days of differentiation in mTeSR + DOX.** For each cell type, we show percent differentiated for FACS analysis of canonical markers for non-clonal (turquoise) and mono-clonal (red) cell lines. **a.** Type II Astrocytes **b.** CD8-positive T-cells **c.** Microglia **d.** Regulatory T-cells **e.** Hepatocytes and **f.** B-cells (mesoderm).

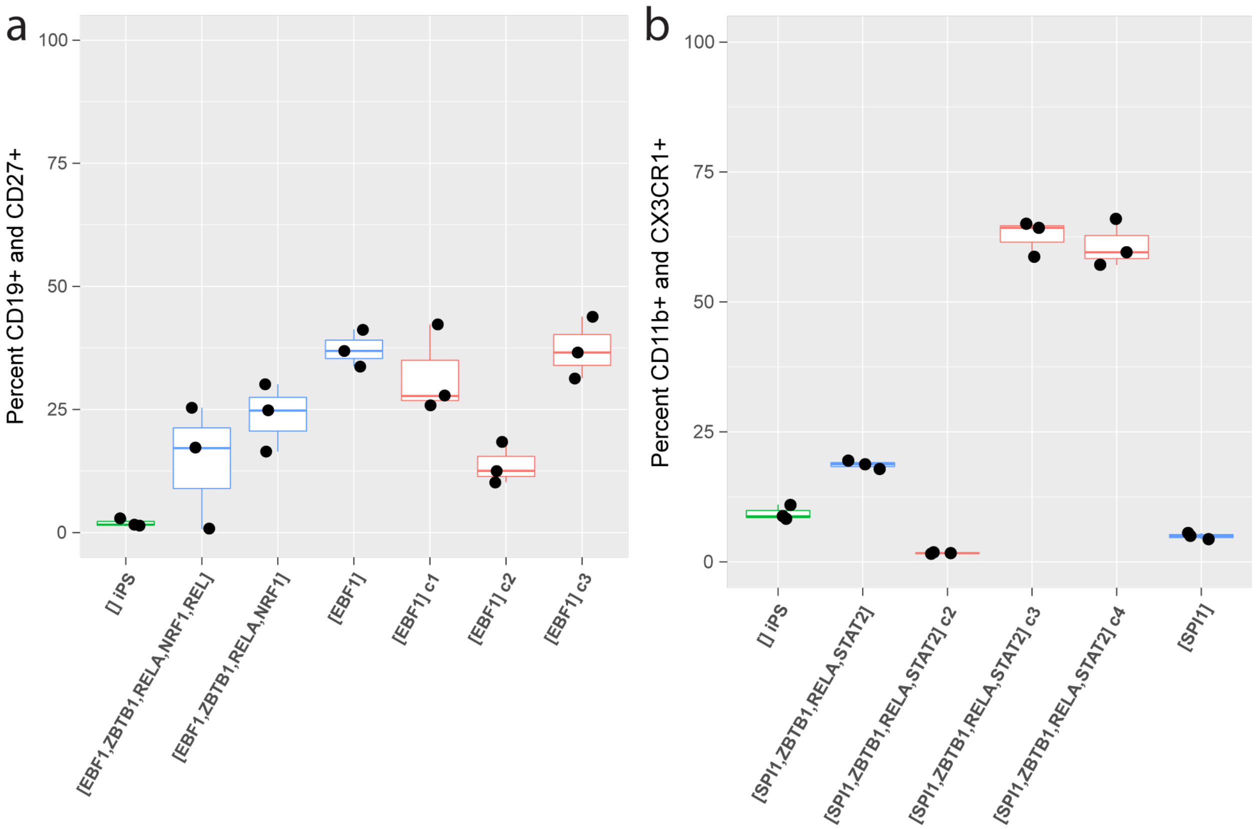


**Supplementary Figure 11: Cell line differentiation for double-positive cell surface markers after 6 days of differentiation in cell-type-specific growth medium + DOX.** For each cell type, we show percent double-positive for FACS analysis of canonical markers for iPSC only (green), non-clonal (turquoise) and mono-clonal (red) cell lines. **a.** B-cells **b.** Microglia

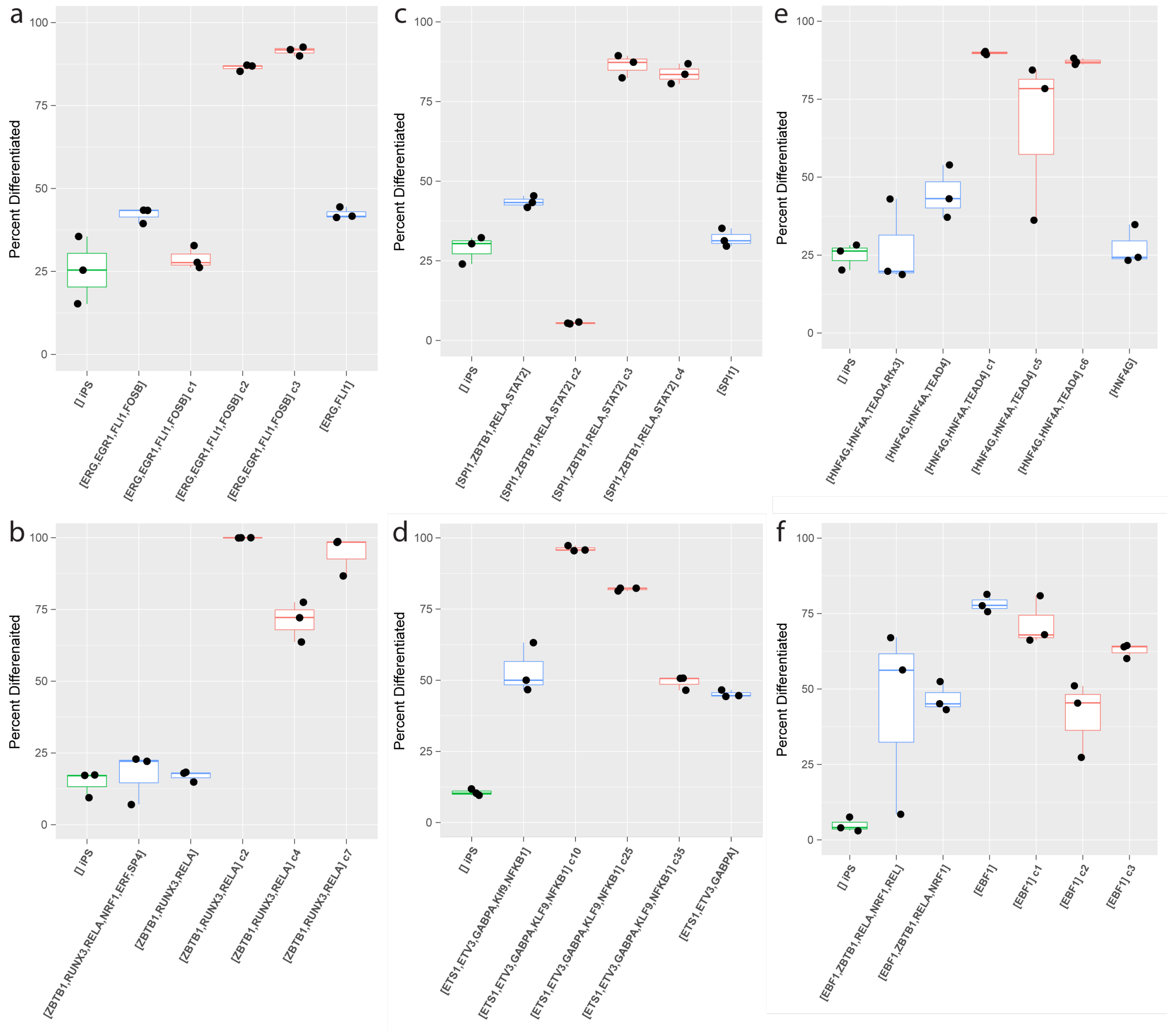


**Supplementary Figure 12: Cell line differentiation for either one or both cell surface markers after 6 days of differentiation in cell-type-specific growth medium + DOX.** For each cell type, we show percent differentiated for FACS analysis of canonical markers for non-clonal (turquoise), mono-clonal (red) cell lines, and an iPSC + media control (green). **a.** Type II Astrocytes **b.** CD8-positive T-cells **c.** Microglia **d.** Regulatory T-cells **e.** Hepatocytes and **f.** B-cells (mesoderm).

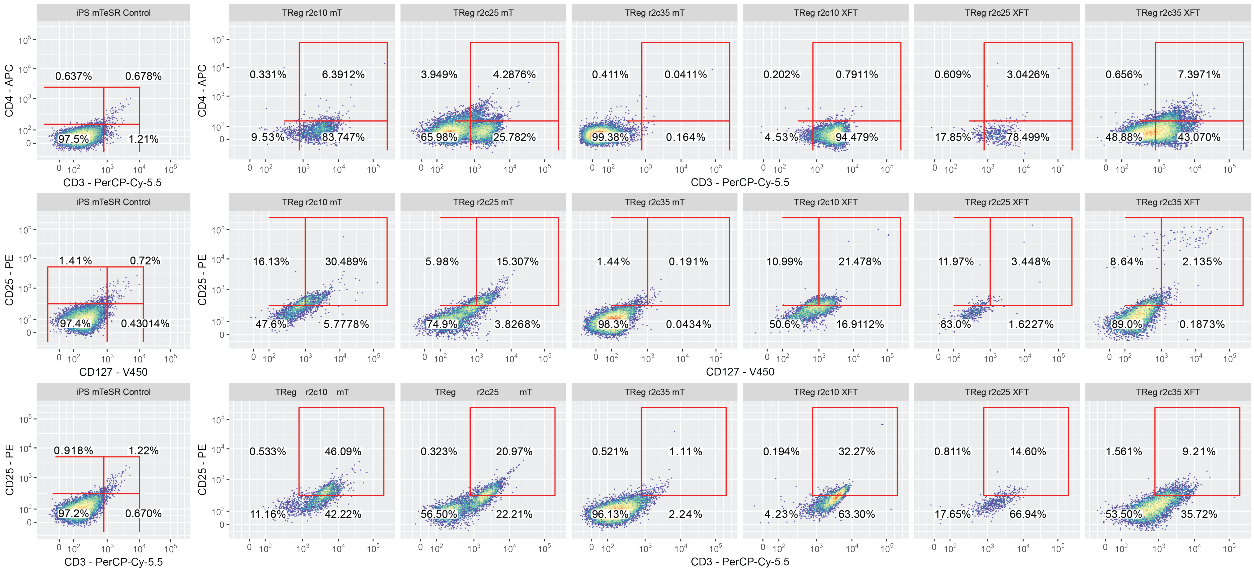


**Supplementary Figure 13:** Additional markers for characterized regulatory T-cell clones in two simple media conditions.

| Tool | Input | | Output | | Can apply to new cell type | Builds Networks | Uses accessible regions | Uses accessibility read counts | Ranks all TFs (supports screnning) | Predicts accesibility | Supports iterative refinement | Biological implementation details considered | Reports new pioneer factor combination discovery | Assumptions |
|---|---|---|---|---|---|---|---|---|---|---|---|---|---|---|
| | RNA-seq | ATAC-seq | TF | PWM | | | | | | | | | | |
| CellNet [3] | x | | x | | | x | | | | | x | | x | Most important TFs will regulate genes within learned biological network |
| Mogrify [4] | x | | x | | | x | | | | | | | x | |
| IRENE [5] | x | x | x | | | x | x | | | | | | x | A core set of regulatory TFs governs cell type identity, conversion efficiency |
| EBseq [6] | x | | x | | x | | | | | | | | | Most important TFs are differentially expressed |
| GarNet [7] | x | x | x | | x | x | x | | x | | | | | Most important TFs regulate differentially expressed genes that contain a nearby motif |
| HOMER [8] | | x | x | x | x | | x | | | | | | | |
| DREME [9] | | x | | x | x | | | | | | | | | Most important TF motifs will be the most over-represented in accessible DNA |
| KMAC [10] | | x | | x | x | | x | | | | | | | |
| AME [11] | | x | x | | x | | x | | | | | | | |
| DeepAccess [12] | | x | x | | x | | x | | x | x | | | | |
| TEPIC [14] | x | x | x | x | x | | x | | | | | | | |
| CellCartographer | x | x | x | x | x | | x | | x | x | x | x | x | |
| diffTF [13] | | x | x | | x | | x | x | x | | | | | Most important TF motifs will be differentially accessible |
| ANANSE [15] | x | x | x | | x | x | x | | | x | | | | |

**Supplementary Table 1: Comparison of tools to identify TFs for cell-type conversion.** Tools for selecting TFs [3, 4, 5, 6, 7, 8, 9, 10, 11, 12, 13, 14, 15] compared for functionality based on [16]

| TOOL | B-cell | T-cell | Macrophage | Mammary Epithelial Cell | Astrocyte | Hepatocyte (Liver) |
|---|---|---|---|---|---|---|
| **Mogrify** | EP300<br>EBF1<br>BACH2<br>PAX5<br>MEF2C<br>SPIB<br>FOS<br>NR3C1 | N/A | MITF<br>SPI1<br>CEBPA<br>MAFB<br>FOSL2<br>IRF1<br>MEF2A<br>SNAI3 | IL1B<br>TP63<br>FOSL1<br>MYC<br>NFKB1<br>HES1<br>JUN<br>FOXA2 | PAX6<br>POU3F2<br>SNAI2<br>RUNX2<br>SOX5<br>E2F5<br>HMGB2<br>SMAD1 | ONECUT1<br>HNF4A<br>NR1H4<br>MLXIPL<br>FOX2A<br>NR5A2<br>ATF5<br>RXRA |
| **IRENE (High Efficiency)** | ETS1<br>FLI1<br>TBX21<br>EBF1<br>IRF4<br>BACH2<br>IKZF1 | N/A | N/A | GRHL3<br>NFYC<br>VDR<br>KLF5<br>MAX | N/A | N/A |
| **CellNet** | PAX5<br>POU2AF1<br>BACH2<br>IRF4<br>SPIB<br>EBF1<br>SP140<br>AFF3<br>ZNF107<br>IKZF3<br>IFT57<br>ZHX2<br>MTERF4<br>ZBTB24<br>ZNF92<br>HSF5<br>IKZF2<br>BCL11A<br>ZNF860<br>ZBTB5<br>ZNF549<br>E2F5<br>TAF1A<br>ZNF572<br>ZIK1<br>ZNF670<br>ZNF239 | TCF7<br>GATA3<br>BCL11B<br>RORA<br>LEF1<br>TXK<br>SCML4<br>ZNF831<br>GF1<br>FOXP3<br>ZNF80<br>RORC | SPI1<br>MAFB<br>CEBPB<br>ZNF438<br>NR1H3<br>NSC<br>HLX<br>ATF5<br>SPHK1<br>BCL3<br>RELB<br>PPARD<br>ZC3H12A<br>GLMP<br>RFX8<br>MITF<br>TBPL2<br>ZBTB17<br>BHLHE41<br>TFPT<br>ETV5 | N/A | N/A | N/A |

**Supplementary Table 2:** Comparison of outputs from tools with publications demonstrating validated novel TF combinations for differentiation [3, 4, 5]. These tools do not allow for the user to readily evaluate new cell types, so microglia, osteoblast, bronchial epithelial cell, kidney epithelial cell, regulatory T-cell, and delta-gamma T-cell predictions are not available from these tools.

| THRESHOLD | MOTIF | GENE |
|---|---|---|
| 1 RPKM | Runt related factors 2 | RUNX3 |
| | EBF1 | EBF1 |
| | Gmeb1 | GMEB1 |
| | Three zinc finger Kruppel related factors 3 | SP4 |
| | KLF9 | KLF9 |
| | Ets related factors 3 | SPI1 |
| | NRF1 | NRF1 |
| | RFX related factors 1 | RFX1 |
| | RFX related factors 1 | MXI1 |
| | RFX related factors 1 | RFX3 |
| | RFX related factors 1 | RFX5 |
| | ZBTB1 | ZBTB1 |
| | NFYB | NFYB |
| | ETV6 | ETV6 |
| 4 RPKM | Runt related factors 2 | RUNX3 |
| | EBF1 | EBF1 |
| | Gmeb1 | GMEB1 |
| | Three zinc finger Kruppel related factors 3 | SP4 |
| | KLF9 | KLF9 |
| | Ets related factors 3 | SPI1 |
| | NRF1 | NRF1 |
| | RFX related factors 1 | RFX1 |
| | RFX related factors 1 | MXI1 |
| | RFX related factors 1 | RFX3 |
| | RFX related factors 1 | RFX5 |
| | ZBTB1 | ZBTB1 |
| | ETV6 | ETV6 |
| | Rel 1 | RELA |
| | Rel 1 | REL |

**Supplementary Table 3:** Transcription factors and corresponding DNA motifs selected for B-cell pooled screen

| THRESHOLD | MOTIF | GENE |
|---|---|---|
| 1 RPKM | Runt related factors 2 | RUNX3 |
| | Runt related factors 2 | RUNX2 |
| | Ets related factors 2 | ETV3 |
| | Ets related factors 2 | ERF |
| | Ets related factors 2 | FLI1 |
| | Ets related factors 2 | GABPA |
| | Ets related factors 2 | ETS1 |
| | Ets related factors 2 | ELK4 |
| | Three zinc finger Kruppel related factors 3 | SP4 |
| | Gmeb1 | GMEB1 |
| | KLF9 | KLF9 |
| | ZBTB1 | ZBTB1 |
| | NRF1 | NRF1 |
| | Rel 1 | REL |
| | NFYB | NFYB |
| 4 RPKM | Runt related factors 2 | RUNX3 |
| | Runt related factors 2 | RUNX2 |
| | Ets related factors 2 | ETV3 |
| | Ets related factors 2 | ERF |
| | Ets related factors 2 | FLI1 |
| | Ets related factors 2 | GABPA |
| | Ets related factors 2 | ETS1 |
| | Ets related factors 2 | ELK4 |
| | Three zinc finger Kruppel related factors 3 | SP4 |
| | Gmeb1 | GMEB1 |
| | KLF9 | KLF9 |
| | ZBTB1 | ZBTB1 |
| | NRF1 | NRF1 |
| | Rel 1 | RELA |
| | Rel 1 | REL |
| | NFYB | NFYB |

**Supplementary Table 4:** Transcription factors and corresponding DNA motifs selected for cytotoxic T-cell pooled screen

| THRESHOLD | MOTIF | GENE |
|---|---|---|
| 1 RPKM | NFYB | NFYB |
| | Ets related factors 2 | ETV5 |
| | Ets related factors 2 | ERF |
| | Ets related factors 2 | GABPA |
| | Ets related factors 2 | ETS1 |
| | Ets related factors 2 | ETV3 |
| | Ets related factors 2 | FLI1 |
| | Ets related factors 2 | ELK4 |
| | Ets related factors 2 | ELK1 |
| | Gmeb1 | GMEB1 |
| | KLF9 | KLF9 |
| | Runt related factors 2 | RUNX3 |
| | Runt related factors 2 | RUNX2 |
| | NF kappaB related factors 1 | NFKB1 |
| 4 RPKM | NFYB | NFYB |
| | Ets related factors 2 | ERF |
| | Ets related factors 2 | GABPA |
| | Ets related factors 2 | ETS1 |
| | Ets related factors 2 | ETV3 |
| | Ets related factors 2 | FLI1 |
| | Ets related factors 2 | ELK4 |
| | Ets related factors 2 | ELK1 |
| | Gmeb1 | GMEB1 |
| | ZKSCAN1 | ZKSCAN1 |
| | Three zinc finger Kruppel related factors 3 | SP4 |
| | Three zinc finger Kruppel related factors 3 | KLF14 |
| | Runt related factors 2 | RUNX3 |
| | NF kappaB related factors 1 | NFKB1 |
| | NF kappaB related factors 1 | NFKB2 |

**Supplementary Table 5:** Transcription factors and corresponding DNA motifs selected for regulatory T-cell pooled screen

**Supplementary Table 6:** Transcription factors and corresponding DNA motifs selected for astrocyte pooled screen

| THRESHOLD | MOTIF | GENE |
|---|---|---|
| 1 RPKM | Hnf4a | HNF4A |
| | Jun related factors 2 | FOSL2 |
| | Jun related factors 2 | FOSL1 |
| | HNF4G | HNF4G |
| | RFX related factors 1 | RFX5 |
| | RFX related factors 1 | MXI1 |
| | RFX related factors 1 | RFX3 |
| | NFIA | NFIA |
| | POU domain factors 1 | HNF1A |
| | Three zinc finger Kruppel related factors 4 | KLF5 |
| | CP2 related factors 1 | GRHL1 |
| | CP2 related factors 1 | TFCP2 |
| | MEIS1 | MEIS1 |
| | TEF 1 related factors 1 | TEAD3 |
| | TEF 1 related factors 1 | TEAD4 |
| | TEF 1 related factors 1 | TEAD1 |
| 4 RPKM | Hnf4a | HNF4A |
| | Jun related factors 2 | FOSL1 |
| | RFX related factors 1 | RFX5 |
| | RFX related factors 1 | MXI1 |
| | POU domain factors 1 | HNF1A |
| | Three zinc finger Kruppel related factors 4 | KLF5 |
| | Three zinc finger Kruppel related factors 4 | KLF4 |
| | MEIS1 | MEIS1 |
| | TEF 1 related factors 1 | TEAD3 |
| | Forkhead 2 | FOXO3 |
| | Forkhead 2 | FOXO4 |
| | CEBP related 2 | DBP |

**Supplementary Table 7:** Transcription factors and corresponding DNA motifs selected for hepatocyte pooled screen

| THRESHOLD | MOTIF | GENE |
|---|---|---|
| 1 RPKM | Ets related factors 3 | SPI1 |
| | Regulators of differentiation 1 | MEF2A |
| | Regulators of differentiation 1 | MEF2C |
| | Regulators of differentiation 1 | MEF2D |
| | Ets related factors 1 | ELF4 |
| | Ets related factors 1 | ELF1 |
| | STAT1::STAT2 | STAT2 |
| | bHLH ZIP factors 5 | USF1 |
| | bHLH ZIP factors 5 | USF2 |
| | NF kappaB related factors 1 | NFKB1 |
| | NF kappaB related factors 1 | NFKB2 |
| | IRF1 | IRF1 |
| | Fos related factors 1 | NFE2 |
| | Rel 1 | RELA |
| | CEBP related 1 | CEBPG |
| 4 RPKM | Ets related factors 3 | SPI1 |
| | Regulators of differentiation 1 | MEF2A |
| | Regulators of differentiation 1 | MEF2C |
| | Regulators of differentiation 1 | MEF2D |
| | Ets related factors 1 | ELF4 |
| | STAT1::STAT2 | STAT2 |
| | bHLH ZIP factors 5 | USF1 |
| | bHLH ZIP factors 5 | USF2 |
| | NF kappaB related factors 1 | NFKB1 |
| | NF kappaB related factors 1 | NFKB2 |
| | IRF1 | IRF1 |
| | Rel 1 | RELA |
| | RFX related factors 1 | MXI1 |
| | RFX related factors 1 | RFX1 |
| | BACH1 var4 | BACH1 |

**Supplementary Table 8:** Transcription factors and corresponding DNA motifs selected for microglia pooled screen

| THRESHOLD | MOTIF | GENE |
| --- | --- | --- |
| 1 RPKM | Jun related factors 2 | JUN |
| | Jun related factors 2 | JUND |
| | TEF 1 related factors 1 | TEAD3 |
| | TEF 1 related factors 1 | TEAD1 |
| | Fos related factors 1 | JDP2 |
| | CP2 related factors 1 | GRHL1 |
| | CP2 related factors 1 | TFCP2 |
| | Runt related factors 2 | RUNX2 |
| | Atf3 | ATF3 |
| | NFIA | NFIA |
| | CEBPA | CEBPA |
| | CEBP related 1 | CEBPD |
| | CEBP related 1 | CEBPB |
| | CEBP related 1 | CEBPG |
| | Three zinc finger Kruppel related factors 4 | KLF4 |
| | Three zinc finger Kruppel related factors 4 | KLF5 |
| 4 RPKM | Jun related factors 2 | JUN |
| | Jun related factors 2 | JUND |
| | TEF 1 related factors 1 | TEAD3 |
| | TEF 1 related factors 1 | TEAD1 |
| | Fos related factors 1 | JDP2 |
| | CP2 related factors 1 | GRHL1 |
| | Atf3 | ATF3 |
| | CEBPA | CEBPA |
| | CEBP related 1 | CEBPD |
| | CEBP related 1 | CEBPB |
| | CEBP related 1 | CEBPG |
| | Three zinc finger Kruppel related factors 4 | KLF4 |
| | Three zinc finger Kruppel related factors 4 | KLF5 |
| | THAP1 | THAP1 |
| | NRF1 | NRF1 |
| | Ets related factors 2 | ERF |

**Supplementary Table 9:** Transcription factors and corresponding DNA motifs selected for bronchial epithelial cell pooled screen

| THRESHOLD | MOTIF | GENE |
|---|---|---|
| 1 RPKM | Runt related factors 2 | RUNX3 |
| | Runt related factors 2 | RUNX2 |
| | Ets related factors 2 | ERF |
| | Ets related factors 2 | ELK1 |
| | Ets related factors 2 | FLI1 |
| | Ets related factors 2 | ETV5 |
| | Ets related factors 2 | ETV3 |
| | Ets related factors 2 | ELK4 |
| | Ets related factors 2 | GABPA |
| | Ets related factors 2 | ETS1 |
| | TCF 7 related factors 2 | TCF7L2 |
| | TCF 7 related factors 2 | LEF1 |
| | NFYB | NFYB |
| | Three zinc finger Kruppel related factors 4 | KLF4 |
| 4 RPKM | Runt related factors 2 | RUNX3 |
| | Runt related factors 2 | RUNX2 |
| | Ets related factors 2 | ERF |
| | Ets related factors 2 | ELK1 |
| | Ets related factors 2 | FLI1 |
| | Ets related factors 2 | ETV3 |
| | Ets related factors 2 | ELK4 |
| | Ets related factors 2 | GABPA |
| | Ets related factors 2 | ETS1 |
| | TCF 7 related factors 2 | LEF1 |
| | NFYB | NFYB |
| | NFYA | NFYA |
| | SP1 | SP1 |
| | ZKSCAN1 | ZKSCAN1 |

**Supplementary Table 10:** Transcription factors and corresponding DNA motifs selected for gamma delta T-cell pooled screen

| THRESHOLD | MOTIF | GENE |
|---|---|---|
| 1 RPKM | Jun related factors 2 | JUND |
| | Jun related factors 2 | JUN |
| | Runt related factors 2 | RUNX2 |
| | POU domain factors 1 | HNF1A |
| | Fos related factors 1 | JDP2 |
| | bZIP 3 | BACH1 |
| | bZIP 3 | BACH2 |
| | Atf3 | ATF3 |
| | RFX related factors 1 | MXI1 |
| | RFX related factors 1 | RFX3 |
| | RFX related factors 1 | RFX1 |
| | RFX related factors 1 | RFX5 |
| | Three zinc finger Kruppel related factors 3 | SP4 |
| 4 RPKM | Jun related factors 2 | JUND |
| | Jun related factors 2 | JUN |
| | Runt related factors 2 | RUNX2 |
| | Fos related factors 1 | JDP2 |
| | Atf3 | ATF3 |
| | RFX related factors 1 | MXI1 |
| | RFX related factors 1 | RFX5 |
| | HMBOX1 | HMBOX1 |
| | REST | REST |
| | Ets related factors 2 | GABPA |
| | Ets related factors 2 | ETS1 |
| | Ets related factors 2 | ERF |
| | Ets related factors 2 | ETV3 |
| | Ets related factors 2 | ELK4 |

**Supplementary Table 11:** Transcription factors and corresponding DNA motifs selected for kidney epithelial cell pooled screen

| THRESHOLD | MOTIF | GENE |
|---|---|---|
| 1 RPKM | Ets related factors 3 | SPI1 |
| | CEBPA | CEBPA |
| | Jun related factors 2 | FOSL2 |
| | Jun related factors 2 | FOS |
| | Jun related factors 2 | JUND |
| | Jun related factors 2 | JUNB |
| | Jun related factors 2 | JUN |
| | Ets related factors 1 | ELF3 |
| | Ets related factors 1 | ELF1 |
| | Ets related factors 1 | ELF4 |
| | ETV6 | ETV6 |
| | Fos related factors 1 | NFE2 |
| | Fos related factors 1 | JDP2 |
| | Three zinc finger Kruppel related factors 3 | SP4 |
| | Three zinc finger Kruppel related factors 4 | KLF4 |
| 4 RPKM | Ets related factors 3 | SPI1 |
| | CEBPA | CEBPA |
| | Jun related factors 2 | FOSL2 |
| | Jun related factors 2 | FOS |
| | Jun related factors 2 | JUND |
| | Jun related factors 2 | JUNB |
| | Jun related factors 2 | JUN |
| | Ets related factors 1 | ELF3 |
| | Ets related factors 1 | ELF1 |
| | Ets related factors 1 | ELF4 |
| | ETV6 | ETV6 |
| | Fos related factors 1 | JDP2 |
| | Three zinc finger Kruppel related factors 4 | KLF4 |
| | Regulators of differentiation 1 | MEF2C |
| | Regulators of differentiation 1 | MEF2A |

**Supplementary Table 12:** Transcription factors and corresponding DNA motifs selected for macrophage pooled screen

| THRESHOLD | MOTIF | GENE |
|---|---|---|
| 1 RPKM | Jun related factors 2 | JUND |
| | Jun related factors 2 | FOSL2 |
| | Jun related factors 2 | FOSL1 |
| | Jun related factors 2 | FOS |
| | Jun related factors 2 | JUNB |
| | Jun related factors 2 | JUN |
| | bZIP 3 | BACH1 |
| | TEF 1 related factors 1 | TEAD3 |
| | TEF 1 related factors 1 | TEAD1 |
| | TEF 1 related factors 1 | TEAD4 |
| | NFIA | NFIA |
| | Fos related factors 1 | JDP2 |
| | Three zinc finger Kruppel related factors 3 | SP4 |
| 4 RPKM | Jun related factors 2 | JUND |
| | Jun related factors 2 | FOSL2 |
| | Jun related factors 2 | FOSL1 |
| | Jun related factors 2 | FOS |
| | Jun related factors 2 | JUNB |
| | Jun related factors 2 | JUN |
| | TEF 1 related factors 1 | TEAD3 |
| | TEF 1 related factors 1 | TEAD1 |
| | TEF 1 related factors 1 | TEAD4 |
| | NFIA | NFIA |
| | CP2 related factors 1 | TFCP2 |
| | THAP1 | THAP1 |
| | Ets related factors 2 | ETS1 |
| | Ets related factors 2 | ERF |

**Supplementary Table 13:** Transcription factors and corresponding DNA motifs selected for mammary epithelial cell pooled screen

| THRESHOLD | MOTIF | GENE |
|---|---|---|
| 1 RPKM | NFIA | NFIA |
| | EBF1 | EBF1 |
| | Bcl6 | BCL6 |
| | Interferon regulatory factors 1 | IRF9 |
| | CDC5L | CDC5L |
| | Nuclear factor 1 1 | NFIC |
| | Nuclear factor 1 1 | NFIX |
| | HMBOX1 | HMBOX1 |
| | TBrain related factors 2 | POU6F1 |
| | SMAD1 | SMAD1 |
| | bZIP 1 | ATF2 |
| | bZIP 1 | GMEB2 |
| | NFAT related factors 1 | NFATC4 |
| | ELF2 | ELF2 |
| | SIX5 var2 | SIX5 |
| 4 RPKM | EBF1 | EBF1 |
| | Bcl6 | BCL6 |
| | Interferon regulatory factors 1 | IRF9 |
| | CDC5L | CDC5L |
| | Nuclear factor 1 1 | NFIC |
| | Nuclear factor 1 1 | NFIX |
| | HMBOX1 | HMBOX1 |
| | SMAD1 | SMAD1 |
| | bZIP 1 | ATF2 |
| | NFAT related factors 1 | NFATC4 |
| | ELF2 | ELF2 |
| | RXR related receptors NR2 2 | RARA |
| | RXR related receptors NR2 2 | RXRA |
| | MZF1 | MZF1 |

**Supplementary Table 14:** Transcription factors and corresponding DNA motifs selected for osteoblast pooled screen

| CELL TYPE | ENCODE IDS | | | |
|---|---|---|---|---|
| common myeloid progenitor CD34-positive | ENCFF644NDH | ENCFF919ARA | ENCFF968XRL | |
| B-cell | ENCFF000ESM | ENCFF209TGG | ENCFF652AIR | |
| CD8-positive alpha-beta T-cell | ENCFF158OUQ | ENCFF214MGP | ENCFF442UGX | |
| foreskin keratinocyte | ENCFF023DDP | ENCFF256VDB | ENCFF676DZK | |
| CD14-positive monocyte | ENCFF000HUU | ENCFF246VSY | ENCFF709TXQ | |
| CD4-positive alpha-beta T-cell | ENCFF141XOX | ENCFF249TVM | ENCFF861MLB | |
| T-cell | ENCFF041PMF | ENCFF429CDV | ENCFF684RDZ | |
| mammary epithelial cell | ENCFF000GDZ | ENCFF014CMH | ENCFF214OUU | |
| trophoblast cell | ENCFF143KXV | ENCFF364YYU | ENCFF524LEU | |
| fibroblast of lung | ENCFF000IJZ | ENCFF000IKA | ENCFF000IKH | |
| fibroblast of skin abdomen | ENCFF079EAZ | ENCFF224MVX | | |
| natural killer cell | ENCFF407PLB | ENCFF619DRO | | |
| skeletal muscle myoblast | ENCFF000GPC | ENCFF000GPO | ENCFF000GPP | |
| endodermal cell | ENCFF023WJA | ENCFF103SIG | ENCFF409SDV | (in vitro differentiated) |
| cardiac muscle cell | ENCFF306XSY | ENCFF335BVH | ENCFF852LNM | (in vitro differentiated) |
| hepatocyte | ENCFF385OMY | ENCFF795SKU | ENCFF987HYN | (in vitro differentiated) |
| myocyte | ENCFF000DZT | ENCFF000DZV | ENCFF000DZW | (in vitro differentiated) |
| astrocyte | ENCFF253MPW | ENCFF393GQZ | ENCFF424DEO | (total RNAseq) |
| dermis blood vessel endothelial cell | ENCFF001QZF | ENCFF001QZG | ENCFF001RAP | (total RNAseq) |
| dermis microvascular lymphatic vessel endothelial cell | ENCFF001RAV | ENCFF001RAW | ENCFF001RBD | (total RNAseq) |
| lung microvascular endothelial cell | ENCFF001RBF | ENCFF001RBG | ENCFF001RBV | (total RNAseq) |
| bronchial epithelial cell | ENCFF001RAB | ENCFF001RAC | ENCFF001RBN | (total RNAseq) |
| epithelial cell of proximal tubule | ENCFF479JTM | ENCFF541NUM | ENCFF886ZUJ | (total RNAseq) |
| fibroblast of dermis | ENCFF000HWI | ENCFF000HXA | ENCFF000HXB | (total RNAseq) |
| fibroblast of villous mesenchyme | ENCFF000GXC | ENCFF000GXU | ENCFF000GXV | (total RNAseq) |
| glomerular endothelial cell | ENCFF605VBT | ENCFF765PKE | ENCFF867UER | (total RNAseq) |
| kidney epithelial cell | ENCFF109IUU | ENCFF231KEB | ENCFF322VHJ | (total RNAseq) |
| microglia | SRR3319495 | SRR3319496 | | (RNA-seq from mouse) |
| macrophage | C57B16 (UCSD) | C57B16 (UCSD) | | (RNA-seq from mouse) |
| T-regulatory cell | SRR6467059 | SRR6467060 | | (RNA-seq from mouse) |
| GM23338 (PGP1) | ENCFF267VJG | ENCFF563KDS | ENCFF736FIO | |
| gdT-cell | SRR8194737 | SRR8194738 | SRR8194739 | |
| microglia (kuppfer) | SRR10067714 | SRR10067715 | | (RNA-seq from mouse) |
| microglia (alveolar) | SRR10084463 | SRR10084464 | | (RNA-seq from mouse) |
| **TISSUE TYPE** | **ENCODE IDS** | | | |
| muscle of arm | ENCFF547NQV | ENCFF995KMZ | | |
| stomach | ENCFF529WDB | ENCFF855FUM | ENCFF952LQK | |
| muscle of back | ENCFF467CIE | ENCFF713VWJ | ENCFF972LVN | |
| small intestine | ENCFF587VJD | ENCFF700QQQ | ENCFF796HDN | |
| muscle of leg | ENCFF038OLY | ENCFF591YBG | ENCFF726EJA | |
| large intestine | ENCFF062QDJ | ENCFF064GDU | ENCFF855WFH | |
| heart | ENCFF076IRZ | ENCFF092JPL | ENCFF975AUW | |
| adrenal gland | ENCFF080VMX | ENCFF345CEU | ENCFF396DVW | |
| kidney | ENCFF390GZH | ENCFF391AJE | ENCFF715EGE | |
| left lung | ENCFF565ABG | ENCFF836QCP | ENCFF994THJ | |
| brain | ENCFF456MMS | ENCFF850ZLY | ENCFF897IUQ | |
| lung | ENCFF136TCV | ENCFF477LPQ | ENCFF704QZC | |
| thymus | ENCFF410KEZ | ENCFF822QPE | ENCFF974RKT | |
| right lung | ENCFF106TWX | ENCFF240TXD | ENCFF939HWL | |
| renal cortex interstitium | ENCFF051TUA | ENCFF149FEQ | ENCFF371QWQ | |
| placenta | ENCFF394BZF | ENCFF494AAG | ENCFF689QAM | |
| spinal cord | ENCFF123FVX | ENCFF659JRI | ENCFF725KJB | |
| left renal cortex interstitium | ENCFF139JYD | ENCFF832ENU | ENCFF859IDO | |
| left renal pelvis | ENCFF453MGJ | ENCFF490TMA | ENCFF493PDU | |
| sigmoid colon | ENCFF232KGN | ENCFF831HCD | ENCFF992HBZ | |
| liver | ENCFF367XVO | ENCFF688YVP | ENCFF938MAS | |
| ovary | ENCFF102XVK | ENCFF254IHO | ENCFF715JXK | |
| right renal cortex interstitium | ENCFF589SOA | ENCFF701WAW | ENCFF711XVI | |
| testis | ENCFF016TGP | ENCFF276UME | ENCFF604DIX | |
| heart left ventricle | ENCFF283NRH | ENCFF299ILO | ENCFF862EDE | |
| mesendoderm | ENCFF226ZUY | ENCFF667BAN | ENCFF771WAJ | (in vitro differentiated) |
| muscle of trunk | ENCFF372ISN | ENCFF703HYW | ENCFF967REG | |
| pancreas | ENCFF240UXB | ENCFF260PST | ENCFF995ZHM | |
| psoas muscle | ENCFF096YUH | ENCFF186JWV | ENCFF830BRN | |
| heart right ventricle | ENCFF360MFU | ENCFF446BWK | ENCFF450DDH | |

**Supplementary Table 15:** Data IDs for transcriptomics data used from ENCODE and GEO in this study. Data is polyA-RNAseq from human primary cells unless otherwise noted.

| CELL TYPE | ENCODE IDS | | | |
|---|---|---|---|---|
| common myeloid progenitor CD34-positive | ENCFF270ZPP | ENCFF331UYM | ENCFF467ADD | |
| B-cell | ENCFF156JWD | ENCFF490CTH | ENCFF726OGS | |
| CD8-positive alpha-beta T-cell | ENCFF093PEA | ENCFF421KOH | ENCFF999HYW | |
| foreskin keratinocyte | ENCFF457CHS | ENCFF537QTE | ENCFF951RBN | |
| CD14-positive monocyte | ENCFF453ORV | ENCFF676UYS | ENCFF960OEQ | |
| CD4-positive alpha-beta T-cell | ENCFF513JFC | ENCFF645BIK | ENCFF772XKG | |
| T-cell | ENCFF275UFF | ENCFF688GBU | ENCFF756BCF | |
| mammary epithelial cell | ENCFF040XDV | ENCFF426PKC | ENCFF914GIZ | |
| trophoblast cell | ENCFF442OAX | ENCFF883LZW | ENCFF917VQR | |
| fibroblast of lung | ENCFF001CBY | ENCFF001CBZ | ENCFF001EDJ | |
| fibroblast of skin abdomen | ENCFF491YLY | ENCFF707PZQ | ENCFF972ZVY | |
| natural killer cell | ENCFF431SXN | ENCFF505OFY | ENCFF521XLX | |
| skeletal muscle myoblast | ENCFF001DNY | ENCFF001DNZ | ENCFF689CAW | |
| endodermal cell | ENCFF602VMH | ENCFF667SWK | ENCFF754PQR | (in vitro differentiated) |
| cardiac muscle cell | ENCFF081PHP | ENCFF206VSW | ENCFF242LGB | (in vitro differentiated) |
| hepatocyte | ENCFF415CLL | ENCFF699RMR | ENCFF754VPL | (in vitro differentiated) |
| myocyte | ENCFF186RIJ | ENCFF675VWC | ENCFF717GPW | (in vitro differentiated) |
| iPS DF 1911 | ENCFF355EAK | ENCFF468OTK | ENCFF932RNN | |
| foreskin fibroblast | ENCFF099TON | ENCFF324KQA | ENCFF602LUO | |
| foreskin melanocyte | ENCFF335ZRE | ENCFF725IKN | ENCFF848LBN | |
| naïve thymus-derived CD4-positive alpha-beta T-cell | ENCFF000SEV | ENCFF000SEX | ENCFF001CSQ | |
| astrocyte | ENCFF001EBK | ENCFF001EBL | | |
| dermis blood vessel endothelial cell | ENCFF001BIA | ENCFF001DIQ | ENCFF001DIR | |
| dermis microvascular lymphatic vessel endothelial cell | ENCFF001BJJ | ENCFF001BJK | ENCFF001DID | |
| lung microvascular endothelial cell | ENCFF001BGT | ENCFF001BGU | ENCFF001DJK | |
| bronchial epithelial cell | ENCFF001EFR | ENCFF001EFT | | |
| epithelial cell of proximal tubule | ENCFF001EFQ | ENCFF001EFS | | |
| fibroblast of dermis | ENCFF001ECC | ENCFF001ECD | | |
| fibroblast of villous mesenchyme | ENCFF001DON | ENCFF001DOO | | |
| glomerular endothelial cell | ENCFF001DMG | ENCFF001DMH | | |
| kidney epithelial cell | ENCFF001DLX | ENCFF001DLY | | |
| microglia | SRR6351241 | SRR6351242 | SRR6351243 | (ATAC-seq from mouse) |
| macrophage | C57B16 (UCSD) | C57B16 (UCSD) | | (ATAC-seq from mouse) |
| T-regulatory cell | SRR5799558 | SRR5799559 | | (ATAC-seq from mouse) |
| GM23338 (PGP1) | ENCFF015PPK | ENCFF608YCF | ENCFF923CNP | |
| gdT-cell | SRR5799430 | SRR5799431 | | (ATAC-seq from mouse) |
| microglia (kuppfer) | SRR8760492 | SRR8760493 | | (ATAC-seq from mouse) |
| microglia (alveolar) | SRR10084471 | SRR10084472 | | (ATAC-seq from mouse) |
| **TISSUE TYPE** | **ENCODE IDS** | | | |
| muscle of arm | ENCFF547NQV | ENCFF995KMZ | | |
| stomach | ENCFF267EXK | ENCFF952LQK | | |
| muscle of back | ENCFF467CIE | ENCFF713VWJ | | |
| small intestine | ENCFF020BOB | ENCFF534QCZ | ENCFF706CDD | |
| muscle of leg | ENCFF277CXU | ENCFF330RAE | ENCFF729ISP | |
| large intestine | ENCFF090TEE | ENCFF653XPD | ENCFF903LSJ | |
| heart | ENCFF168YBC | ENCFF700IMY | ENCFF976EET | |
| adrenal gland | ENCFF031XLZ | ENCFF650AZH | ENCFF813STM | |
| kidney | ENCFF489SPU | ENCFF640OOB | ENCFF912EKE | |
| left lung | ENCFF055SAI | ENCFF056BQQ | ENCFF668YLU | |
| brain | ENCFF456IBX | ENCFF587MSJ | ENCFF829USQ | |
| lung | ENCFF234IRY | ENCFF852BOV | ENCFF878NNA | |
| thymus | ENCFF583SLL | ENCFF649VDK | ENCFF800OUU | |
| right lung | ENCFF072FIM | ENCFF404ADQ | ENCFF826TDY | |
| renal cortex interstitium | ENCFF171ALU | ENCFF457FVL | ENCFF516MYU | |
| placenta | ENCFF044IHA | ENCFF618VTK | ENCFF700KZI | |
| spinal cord | ENCFF260GSJ | ENCFF354LAX | ENCFF369QCM | |
| left renal cortex interstitium | ENCFF159XPF | ENCFF711KKQ | ENCFF776RFT | |
| left renal pelvis | ENCFF423HOT | ENCFF544DPT | ENCFF946SSJ | |
| sigmoid colon | ENCFF255JSR | ENCFF270DKA | ENCFF333LUA | |
| liver | ENCFF005CBC | ENCFF490OVY | ENCFF727HFJ | |
| ovary | ENCFF353LZV | ENCFF527WZF | ENCFF697AAT | |
| right renal cortex interstitium | ENCFF102SFO | ENCFF391HSJ | ENCFF709HXO | |
| testis | ENCFF048RXA | ENCFF349IJY | ENCFF720UVP | |
| heart left ventricle | ENCFF049ZJG | ENCFF172MDS | ENCFF545EPR | |
| mesendoderm | ENCFF447LVL | ENCFF514CTP | ENCFF572UGZ | (in vitro differentiated) |
| muscle of trunk | ENCFF065TLM | ENCFF258AND | ENCFF516RIM | |
| pancreas | ENCFF648AKE | ENCFF822MKL | ENCFF859WBE | |
| psoas muscle | ENCFF239ACE | ENCFF464GIS | ENCFF802BTC | |
| heart right ventricle | ENCFF431CNR | ENCFF681BJT | ENCFF758SIA | |
| left kidney | ENCFF052VQG | ENCFF443QJN | ENCFF768FPA | |
| spleen | ENCFF569VXR | ENCFF834OBR | ENCFF957FCM | |
| urinary bladder | ENCFF002DZB | ENCFF002DZD | ENCFF002DZE | |

**Supplementary Table 16:** Data IDs for epigenetics data used from ENCODE and GEO in this study. Data is DNAse-seq from human primary cells unless otherwise noted.

| Name | Sequence |
|---|---|
| Barcode_amp1_F | GTGACTGGAGTTCAGACGTGTGCTCTTCCGCAACAGATGGCTGGC |
| Barcode_amp1_R | ACACTCTTTCCCTACACGACGCTCTTCCGATCTTCCAAGCACCTGCTACATAGC |
| Barcode_amp2_F | GTGACTGGAGTTCAGACGTGTGCTCTTCCGAGGAAAGGACAGTGGGAGTGGC |
| Barcode_amp2_R | ACACTCTTTCCCTACACGACGCTCTTCCGATCGCCTTTTCCAAGCACCTGCTACATAGC |
| Ad1.1_TCCACGAA | AATGATACGGCGACCACCGAGATCTACACTCCACGAATCGTCGGCAGCGTCAGATGT |
| Ad1.2_ATCGCTAC | AATGATACGGCGACCACCGAGATCTACACATCGCTACTCGTCGGCAGCGTCAGATGT |
| Ad1.3_GACCTTCA | AATGATACGGCGACCACCGAGATCTACACGACCTTCATCGTCGGCAGCGTCAGATGT |
| Ad2.1_TAAGGCGA | CAAGCAGAAGACGGCATACGAGATTCGCCTTAGTCTCGTGGGCTCGGAGATGT |
| Ad2.2_CGTACTAG | CAAGCAGAAGACGGCATACGAGATCTAGTACGGTCTCGTGGGCTCGGAGATGT |
| Ad2.3_AGGCAGAA | CAAGCAGAAGACGGCATACGAGATTTCTGCCTGTCTCGTGGGCTCGGAGATGT |
| Ad2.4_TCCTGAGC | CAAGCAGAAGACGGCATACGAGATGCTCAGGAGTCTCGTGGGCTCGGAGATGT |
| Ad2.5_GGACTCCT | CAAGCAGAAGACGGCATACGAGATAGGAGTCCGTCTCGTGGGCTCGGAGATGT |
| Ad2.6_TAGGCATG | CAAGCAGAAGACGGCATACGAGATCATGCCTAGTCTCGTGGGCTCGGAGATGT |
| Ad2.7_CTCTCTAC | CAAGCAGAAGACGGCATACGAGATGTAGAGAGGTCTCGTGGGCTCGGAGATGT |
| Ad2.8_CAGAGAGG | CAAGCAGAAGACGGCATACGAGATCCTCTCTGGTCTCGTGGGCTCGGAGATGT |
| Ad2.9_GCTACGCT | CAAGCAGAAGACGGCATACGAGATAGCGTAGCGTCTCGTGGGCTCGGAGATGT |
| Ad2.10_CGAGGCTG | CAAGCAGAAGACGGCATACGAGATCAGCCTCGGTCTCGTGGGCTCGGAGATGT |
| Ad2.11_AAGAGGCA | CAAGCAGAAGACGGCATACGAGATTGCCTCTTGTCTCGTGGGCTCGGAGATGT |
| Ad2.12_GTAGAGGA | CAAGCAGAAGACGGCATACGAGATTCCTCTACGTCTCGTGGGCTCGGAGATGT |
| Ad2.13_GTCGTGAT | CAAGCAGAAGACGGCATACGAGATATCACGACGTCTCGTGGGCTCGGAGATGT |
| Ad2.14_ACCACTGT | CAAGCAGAAGACGGCATACGAGATACAGTGGTGTCTCGTGGGCTCGGAGATGT |
| Ad2.15_TGGATCTG | CAAGCAGAAGACGGCATACGAGATCAGATCCAGTCTCGTGGGCTCGGAGATGT |
| Ad2.16_CCGTTTGT | CAAGCAGAAGACGGCATACGAGATACAAACGGGTCTCGTGGGCTCGGAGATGT |
| Ad2.17_TGCTGGGT | CAAGCAGAAGACGGCATACGAGATACCCAGCAGTCTCGTGGGCTCGGAGATGT |
| Ad2.18_GAGGGGTT | CAAGCAGAAGACGGCATACGAGATAACCCCTCGTCTCGTGGGCTCGGAGATGT |
| Ad2.19_AGGTTGGG | CAAGCAGAAGACGGCATACGAGATCCCAACCTGTCTCGTGGGCTCGGAGATGT |

**Supplementary Table 17:** Oligonucleotids used in this study.
